# Structural and kinetic insights into tRNA promoter engagement by yeast general transcription factor TFIIIC

**DOI:** 10.1101/2024.08.28.610035

**Authors:** Wolfram Seifert-Dávila, Anastasiia Chaban, Florence Baudin, Mathias Girbig, Luis Hauptmann, Thomas Hoffmann, Olivier Duss, Sebastian Eustermann, Christoph W. Müller

**Affiliations:** European Molecular Biology Laboratory (EMBL), Structural and Computational Biology Unit, Meyerhofstrasse 1, 69117 Heidelberg, Germany; Candidate for joint PhD degree from EMBL and Faculty of Biosciences, Heidelberg University, 69120 Heidelberg, Germany; Max Planck Institute for Terrestrial Microbiology, Karl-von-Frisch Straße 10, 35043 Marburg, Germany

## Abstract

Transcription of tRNA genes by RNA polymerase III requires the general transcription factor IIIC (TFIIIC), which recognizes intragenic A-box and B-box DNA motifs of type II gene promoters. However, the underlying mechanism has remained elusive, in part due to missing structural information for A-box recognition. In this study, we use single-particle cryo-EM and single-molecule FRET (smFRET) to reveal structural and real-time kinetic insights into how the 520 kDa yeast TFIIIC complex engages A- and B-box DNA motifs in the context of a tRNA gene promoter. Cryo-EM structures of τA and τB subcomplexes bound to the A- and B-box were obtained at 3.7 and 2.5 Å resolution, respectively, while cryo-EM single particle mapping determined the specific distance and relative orientation of the τA and τB subcomplexes revealing a fully engaged state of TFIIIC. smFRET experiments show that overall recruitment and residence times of TFIIIC on a tRNA gene are primarily governed by B-box recognition, while footprinting experiments suggest a key role of τA and the A-box in TFIIIB and Pol III recruitment following TFIIIC recognition of type II promoters.

## Introduction

Transcription of small non-coding RNAs, such as 5S rRNA, tRNA, 7SL and U6 small nuclear RNA is carried out by RNA polymerase (Pol) III (1,2). Most Pol III genes contain gene-internal promoters regions (type I and type II promoters), although in some Pol III genes the promoter is also located outside the gene (type III promoter) (3). Gene-internal regions recognized by Pol III-specific transcription factors are called internal control regions (ICRs). ICRs are highly conserved and discontinuous DNA segments, which, depending on their consensus sequences, are named A-box, B-box and C-box (4). In the case of tRNA gene promoters (type II), the multi-subunit general transcription factor (TF) IIIC binds to the A-box and B-box, while in 5S rRNA gene promoters (type I), the presence of an A-box, intermediate element (IE) and C-box allows the binding of TFIIIA followed by binding of TFIIIC (5–7). TFIIIC is an assembly factor, because it positions TFIIIB upstream of the transcription start site, which in turn recruits Pol III (8). TFIIIB harbors three subunits: TATA-binding protein (TBP), B double prime (Bdp1), and B-related factor 1 (Brf1) (9). TFIIIC is dispensable once transcription starts, and it is TFIIIB, alone, which positions Pol III for repeated transcription cycles (10).

The interaction of TFIIIC with tRNA gene promoters is predominantly mediated by the B-box, which interacts with TFIIIC with nanomolar (nM) affinity (11), while the A-box exhibits a comparatively weaker micromolar (µM) affinity (9,12). The segment between A- and B-box has been shown to be non-essential for transcription, as point mutations in this area do not significantly alter the transcriptional activity (13). However, the length of this intervening region influences the affinity of TFIIIC for its intragenic promoters (14), with the number of nucleotides between A- and B-box ranging from 31 to 93 bp (15). To adapt to these naturally varying distances, TFIIIC interacts with the A-box and B-box through its two subcomplexes, τA and τB, respectively, which are connected by a flexible linker (16,17).

In a function independent of Pol III transcription, genome-wide studies in several eukaryotes identified TFIIIC at specific genomic regions, called ETC (Extra TFIIIC sites), in the absence of TFIIIB or Pol III (18–20). Many of these sites are in close proximity to binding sites of CCCTC-binding factor (CTCF), a protein involved in forming chromatin boundaries (20). Additionally, TFIIIC has been associated with chromosome architectural proteins such as condensin and cohesin, as well as chromatin remodelers like SMARCD1 (21–23). These findings suggest a general role of TFIIIC as a global genome organizer (24).

In *Saccharomyces cerevisiae*, TFIIIC is a multi-subunit complex of approximately 520 kDa composed of six subunits: τ138, τ131, τ95, τ91, τ60, and τ55, each identified and heterologously cloned (9,15,25–30). Initial attempts to reconstitute the minimal TFIIIC τA and τB subcomplexes divided each module into three subunits each: τ131, τ95, and τ55 for τA and τ138, τ91, and τ60 for τB, where the τB subcomplex exhibited strong DNA-binding capabilities; whereas the τA subcomplex only showed non-specific DNA-binding (31). Recent structural studies of yeast and human TFIIIC bound to type I and type II promoters, respectively, demonstrated that only the N-terminal region of τ138 (TFIIIC220 in human) is part of τB, while the C-terminal region of τ138 is an integral part of τA (32,33).

X-ray crystallography provided first structural insights into domains, subunits and subcomplexes of TFIIIC. The τ60/Δτ91 (residues 159–672) heterodimer was identified as forming the stable core of the τB subcomplex (34). The first structurally characterized domain of τ138 was a conserved extended winged-helix (eWH) domain, one of many winged-helix domains predicted for τ138, offering preliminary insights into its interaction with DNA (35). For the τA subcomplex, the structure of the phosphatase domain of τ55 provided clues about the potential role of TFIIIC in linking transcription and metabolism (36). The *S. pomb*e Sfc1/Sfc7 heterodimer (τ55/τ95 in *S. cerevisiae*), which dimerizes through a triple β-barrel, showed resemblance to the general transcription factor TFIIF, while the putative DNA-binding domain (DBD) of Sfc7 suggested different potential DNA-binding modes (12). The structure of the N-terminal tetratricopeptide repeat (TPR) domain of τ131, containing 10 repeats, offered clues about its role in binding to the multiple subunits of TFIIIB (35). Finally, by combining these crystal structures with cross-linking mass spectrometry, a general model of the molecular architecture of the complete TFIIIC was proposed (35).

More recently, cryo-EM studies of the minimal τA subcomplex of yeast TFIIIC suggested that τA acts as a molecular ruler positioning TFIIIB at a fixed distance upstream of the A-box (9). Furthermore, the structure of the yeast TFIIIA-TFIIIC-Brf1-TBP complex bound to a 5S rRNA gene (type I promoter) showed sharp bending of the DNA wrapping around the TFIIIA-TFIIIC complex supported by Brf1-TBP (33). The structure of the human TFIIIC complex was determined in its unbound state and bound to a tRNA gene, demonstrating how τB recognizes its B-box through shape and DNA sequence readout, while τA is connected by a flexible linker to τB that comprises the middle part of the τ138 subunit (human TFIIIC220) (32).

In the cryo-EM structure of human TFIIIC, τA was only observed in its DNA unbound form. Therefore, it has remained unknown, how τA and τB are arranged relative to each other on type II promoter DNA when bound to their respective A-box and B-box binding sites. Our study addresses this gap by determining cryo-EM structures of the yeast τA and τB subcomplexes bound to A- and B-box, respectively, in their fully engaged state in complex with a tRNA gene promoter. smFRET experiments provide real-time insights into binding kinetics of TFIIIC, showing both short (∼ 1 second) and long (∼ 50 seconds) binding events, and also highlight the dynamic DNA sampling by the structurally not-resolved part of τ95 during long TFIIIC binding events. Our results confirm the importance of the B-box for TFIIIC binding, but also show that the A-box does not significantly contribute to TFIIIC retention on the DNA but likely is responsible for downstream TFIIIB and Pol III recruitment.

## Materials and Methods

### Cloning, expression and purification of recombinant TFIIICΔtail and ybbR-TFIIICΔtail complex

The six insect cell codon-optimized *Saccharomyces cerevisiae* TFIIIC subunits were cloned into the pbiGBac2ab vector using the biGBac assembly method (37). The τ95 subunit was truncated at amino acid 593 to create TFIIICΔtail, a variant that is lacking a C-terminal region auto-inhibiting DNA-binding (9). To assemble the ybbR-TFIIICΔtail variant for smFRET studies, an 11-amino acid ybbR tag (DSLEFIASKLA) was introduced between residues 286 and 287 of τ95. To generate the baculovirus, standard protocols were followed. The TFIIICΔtail complex was expressed in High Five cells, while the ybbR-TFIIICΔtail variant was produced in SF21 cells, both using a 1:1000 virus dilution. Cells were harvested at 90-95% viability, centrifuged, washed with 1x PBS, and stored at -80°C.

For purification, cell pellets from 6 L culture were resuspended in lysis buffer (20 mM HEPES pH 7.5, 500 mM NaCl, 2 mM MgCl_2_, 4 mM β-mercaptoethanol, 10% glycerol), using 3 mL of buffer per gram of cell pellet. Protease inhibitors (1 tablet per 20 g cells, Sigma-Aldrich), Benzonase (4 µL per 50 mL buffer), and DNase I (500 µL, 10 mg/mL) were added. The mixture was stirred on ice, sonicated for 3 min at 40% amplitude, and ultracentrifuged at 35,000 rpm for 1 hour at 4°C. The supernatant was incubated with 2 mL of Strep-Tactin Sepharose™ beads (pre-equilibrated with strep-wash buffer: 20 mM HEPES pH 7.5, 150 mM NaCl, 5 mM DTT, 5% glycerol) for 2 hours at 4°C. Beads were washed with 30 mL strep-wash buffer and eluted with strep-elute buffer (50 mM biotin in strep-wash buffer).

Eluates were further purified using a Capto HiRes Q 5/50 column previously equilibrated with Capto-A buffer (20 mM HEPES pH 7.5, 150 mM NaCl, 5 mM DTT). Elution was performed with a linear gradient from 0 to 70% Capto-B buffer (20 mM HEPES pH 7.5, 1 M NaCl, 5 mM DTT), follow by a step gradient to 100% Capto-B buffer. Fractions with conductivity of 25 to 32 mS/cm were pooled, buffer exchanged into Capto-A buffer, concentrated, aliquoted, and flash-frozen for storage.

### DNA oligonucleotides preparation for cryo-EM studies

Template and non-template oligonucleotides corresponding to the yeast tH(GUG)E2 gene (from now on referred to as ^His^tRNA gene) with and without the upstream TFIIIB binding site, 85bp ^His^tRNA and 120bp ^His^tRNA DNA, respectively, were synthesized by Sigma-Aldrich. In the 120bp ^His^tRNA construct, four nucleotides were mutated to introduce a TATA-like sequence aimed at enhancing TFIIIB stability. Only the non-template strand is depicted. The non-template sequence for 85bp ^His^tRNA DNA reads:

5’-TGAAAAGTC**G**CCATCT<u>TAGTATAGTGG</u>TTAGTACACATCGTTGTGGCCGATGAAA CCCT*GGTTCGATTCT*AGGAGATGGCATTTT-3’. The non-template sequence for the 120bp^His^ tRNA DNA is as follows: 5’-GTATTACTCGAGCCCGTATATAAACAGTTCTCCATTGAAAAGTC**G**CCATCT<u>TAGTATAGTGG</u>TTAGTACACATCGTTGTGGCCGATGAAACCCT*GGTTCGATTCT*AGGAGATGGCATTTT-3’. Modifications include the four nucleotides that replaced the wild type sequence (indicated in red) with the bold nucleotide marking the transcription start site (+1 nucleotide). The A-box is underlined, and the B-box is italicized. Annealing was performed in dH_2_O by denaturation at 95°C for 5 min followed by cooling to 20°C at 1°C per min. Annealed DNA was subjected to size-exclusion chromatography using a Superdex 200 Increase 3.2/300 column (Cytiva), equilibrated with a buffer containing 20 mM HEPES pH 7.5, 150 mM NaCl, 5 mM MgCl_2_, and 5 mM DTT.

### Mass photometry experiments

Coverslips (24 mm x 50 mm) were washed with ddH_2_O and subsequently with isopropanol, dried with compressed air, and fitted with a silicone gasket with six wells. To investigate the influence of ionic strength on the binding stability of the TFIIICΔtail complex to the 85 bp and 120 bp ^His^tRNA gene constructs, 0.4 µM of TFIIIC was mixed with an equivalent molar ratio of tRNA gene DNA oligonucleotides. The TFIIICΔtail-DNA mixture was then applied to a Zeba Spin desalting column (ThermoFisher Scientific) pre-equilibrated with a buffer containing 20 mM HEPES pH 8, 2 mM MgCl_2_, 5 mM DTT, and KCl concentrations varying between 150 mM to 250 mM in 25 mM increments. For mass photometry, 19 μL of buffer and 1 μL of the TFIIICΔtail complex (400 nM) were added to each well, achieving a final concentration of 20 nM.

To assess the concentration-dependent oligomerization of TFIIICΔtail and TFIIICΔtail-DNA complexes, a buffer containing 20 mM HEPES pH 8, 100 mM KCl, 2 mM MgCl_2_, 5 mM DTT was used. Sample volumes were adjusted to achieve final concentrations of 50 nM and 100 nM by using 2.5 µL and 5 µL of the sample, respectively, with corresponding adjustments to the buffer volumes.

Experiments were conducted using a Refeyn TwoMP mass photometer (Refeyn Ltd., Oxford, UK), recording one-minute videos with the AcquireMP software (Refeyn Ltd., version 2.4.0) with an image dimension set to 150 × 59 binned pixels, correlating to an imaging area of 10.9 μm x 4.3 μm and a detection zone of 46.3 μm². Data analysis was performed using DiscoverMP software (Refeyn Ltd., version 2.4.0) with a standard contrast-to-mass calibration curve generated using proteins such as bovine serum albumin and immunoglobulin G.

### Filter binding assay

Oligonucleotides (Sigma-Aldrich, HPLC purified) corresponding to both strands of the tRNA His promoter from positions −45 to +76 or DNA mutants were end labeled using [γ-^32^P] adenosine 5′-triphosphate (ATP) and T4 polynucleotide kinase (New England Biolabs) and purified on a 10% acryl/bisacrylamide, 8.3 M (w/v) urea gel. DNA was eluted overnight from excised gel bands in 0.5 M ammonium acetate, 10 mM magnesium acetate, 0.1% (w/v) SDS, 0.1 mM EDTA, and then ethanol precipitated. The labeled strand was annealed with its cold complementary strand at room temperature for 30 min in 20 mM HEPES pH 7.5, 5 mM MgCl_2_, 100 mM KCl after heat denaturation at 95°C for 3 min. Filter binding assays were performed as follows : DNA (∼30,000 cpm, ∼10 nM) was incubated with increasing amounts of TFIIICΔCtail (0.5 nM to 1 μM) in buffer FB (20 mM HEPES pH 7.5, 150 mM KCl, 2 mM MgCl_2_, 5 mM DTT) for 1 hour at 4°C and then filtered through a 0.45 μm nitrocellulose filter (Whatman), pre-equilibrated in the same buffer. Filters were counted in a Tri-Carb 2800TR Cerenkov scintillation counter (Perkin Elmer). Counts were normalized, and a Hill equation with a fixed Hill coefficient of 1 was fitted using Prism (GraphPad).

### EMSA

The DNA strands were first annealed as described above, and diluted in buffer FB. The EMSA reactions typically contained 100 nM to 1 µM TFIIICΔCtail, 10 nM radioactively labeled DNA (150 fmol, 5 kcpm) and 1 µg poly dI/dC double strand (Amersham), in buffer FB. Reactions were incubated for 20 min at RT before loading onto a native 5% acrylamide gel, and run at 4°C at 100 V for 4 h in Tris glycine buffer. Subsequently, the gel was dried for 1 h at 80°C and the signal visualized by exposure to a phosphorimaging screen.

### Footprinting experiments

Labeled DNA strand was hybridized with its cold complementary DNA strand as described before. DNA (0.1 μM) was incubated with TFIIICΔCtail (0.4 and 0.8 μM), in FB buffer for 30 min at RT. Next, the reaction was supplemented with 2.5 mM MgCl_2_ and 0.5 mM CaCl_2_, and DNase I (New England Biolabs) was added to a final concentration of 0.012 U/μl and the reaction incubated for 5 min at 28 °C. Last, DNA was purified by phenol-chloroform extraction, followed by ethanol precipitation, and resuspended in loading buffer (95% (v/v) formamide, 1× TBE, 0.025% (w/v) xylen cyanol, bromphenol blue). As controls, DNase I was omitted in one reaction, and protein in another. Reactions were heated for 2 min at 95°C and analyzed on a denaturing 12% (w/v) polyacrylamide gel (19:1 acrylamide-bisacrylamide, 8.3 M urea, 1× TBE). The gel was exposed to a phosphorimaging screen (Fujifilm), which was then scanned using a Typhoon FLA 9500 laser scanner (GE Healthcare).

### Sample preparation for TFIIICΔtail-DNA complexes for cryo-EM

Buffer screening and sample preparation for TFIIICΔtail-DNA complexes with and without additional transcription factors were carried out for cryo-EM studies. Previous studies in yeast suggested an optimal salt concentration of 135 mM KCl for TFIIIC-^Glu^tRNA_3_ interaction, with concentrations of 200 mM or higher resulting in complex dissociation as evidenced by the loss of DNase I footprinting (38). Comparison between wild type and mutant yeast TFIIIC revealed an optimal binding affinity to ^Glu^tRNA_3_ at 150 mM KCl in gel shift assays (39). A similar analysis of human TFIIIC showed a lower optimal salt concentration of 70 mM KCl (40).

Mass photometry assessed TFIIICΔtail interaction with the ^His^tRNA gene at various salt concentrations, showing interaction up to 200 mM KCl (Supplementary Figure S2A). These conditions were also tested for cryo-EM sample preparation to optimize particle homogeneity and minimize aggregation (Supplementary Figure S2B).

For complex reconstitution, TFIIICΔtail was prepared at 1.97 µM and mixed with an equimolar amount of the 85 bp ^His^tRNA oligonucleotides. This mixture underwent buffer exchange using a Zeba Spin desalting column, pre-equilibrated with a cryo-EM buffer (20 mM HEPES pH 8.0, 2 mM MgCl_2_, 5 mM DTT), and adjusted to varying KCl concentrations of 175, 200, and 225 mM. Screening of these conditions was performed using a Talos™ Arctica™ microscope, with 200 mM KCl selected for data acquisition on a Titan Krios G3 microscope (dataset 1). To stabilize the TFIIICΔtail-DNA complex with other transcription factors, similar strategies were applied, maintaining equimolar concentrations (1.87 µM) of all components. Testing different buffer was important to facilitate the identification of optimal conditions for data acquisition. The chosen conditions for the four additional datasets are described as follows: Dataset 2 included TFIIIC, 120 bp ^His^tRNA gene, Brf1-TBP fusion protein, Bdp1, and Fpt1 in cryo-EM buffer with 150 mM KCl. Dataset 3 comprised TFIIIC, 120 bp ^His^tRNA gene, and Fpt1 in cryo-EM buffer with 175 mM KCl. Dataset 4 contained TFIIIC, 120 bp ^His^tRNA gene, Brf1-TBP fusion protein, Bdp1, and Fpt1 in cryo-EM buffer with 75 mM KCl. Finally, Dataset 5 included TFIIIC, 120 bp ^His^tRNA gene, and Brf1-TBP fusion protein in cryo-EM buffer with 75 mM KCl.

The grid preparation involved plasma cleaning (10% argon, 90% oxygen) for 2 min and 30 seconds on an Ultrafoil R2/2 Au 200 grid. The vitrobot Mark IV was set to 6°C with 100% humidity. To reduce air-water interface interactions of the applied complexes, octyl-glucoside detergent was added to the samples (0.1% for dataset 1, 0.05% for others) before plunge freezing.

### EM data collection and processing of TFIIICΔtail-DNA complex (dataset 1)

To obtain the structure of the yTFIIICΔtail-DNA complex, dataset 1 was collected, consisting of 19,047 image stacks with 40 frames each. Data collection was performed on a Titan Krios G3 electron microscope (ThermoFischer Scientific) at 300 keV, equipped with an energy filter and a Gatan K3 direct electron detector. Images were taken with a total electron dose of 39.6 electrons per Å² and a defocus range set from 0.7 to 1.7 μm, at a magnification of ×105,000, resulting in an effective pixel size of 0.822 Å.

Pre-processing was conducted using RELION 3.1.3 (41). An initial amount of 1,004,005 particles were picked using WARP (42). 2D classification revealed three distinctive classes: τB-DNA dimer, τB-DNA monomer, and τA-DNA monomers. Initial *ab initio* maps were created from these classes followed by heterogeneous refinement in cryoSPARC (43), incorporating two ’junk’ classes of randomly selected particles (Supplementary Figure S3).

For the τB-DNA dimer class, 199,781 dimer particles were identified and subjected to two rounds of heterogeneous refinement and 2D classification, followed by TOPAZ training and picking, yielding 336,587 new particles. Further sorting through two rounds of heterogeneous refinement led to a map of 122,607 particles, followed by non-uniform refinement in cryoSPARC. However, this map showed a preferred orientation (Supplementary Figure S3, bottom left).

The initial processing of τB-DNA monomer class identified 293,371 particles, which were further classified through two rounds of heterogeneous refinement and 2D classification, followed by TOPAZ training and picking, resulting in a newly set of 589,216 particles. These particles were further processed through two additional rounds of heterogeneous refinement, followed by non-uniform refinement in cryoSPARC. The 180,457 particles contributing to this map were then re-imported into RELION for a second round of TOPAZ training and picking. This process yielded a final set of 546,857 particles, iteratively classified through two rounds of heterogeneous refinement. The resulting τB-DNA monomer map, containing 258,272 particles, achieved a resolution of 3.21 Å after non-uniform refinement in cryoSPARC (Supplementary Figure S3, bottom middle).

For the τA-DNA subcomplex, initial 2D classes were used directly for TOPAZ training and picking, generating 271,890 particles. A specific 2D classification step selected 26,803 particles, used for a second round of TOPAZ training and picking, resulting in 750,337 particles. Using 50,000 random particles to create new ’junk classes,’ and performing 2D classification to select particles for the initial τA-DNA map, heterogeneous refinement selected 125,533 particles, which reached a resolution of 6.54 Å after non-uniform refinement in cryoSPARC (Supplementary Figure S3, bottom right).

### EM data collection and processing of TFIIICΔtail-DNA complex with additional transcriptions factors (dataset 2 to dataset 5)

To improve the cryo-EM reconstruction of the τB-DNA and τA-DNA subcomplex, we collected four additional datasets (dataset 2-5), adding extra transcription factors during sample preparation, though these factors were not detected in the final cryo-EM densities (Supplementary Figure S4 and S5). Data were collected on a Titan Krios G3 electron microscope (ThermoFischer Scientific), using the same specifications as dataset 1. Specifically for dataset 2, 16,289 micrographs with a total electron dose of 43.6 electrons/Å² were collected; for dataset 3, 11,644 micrographs at 43.2 electrons/Å²; for dataset 4, 16,169 micrographs at 43.2 electrons/Å²; and for dataset 5, 15,614 micrographs were collected at a dose of 44.4 electrons/Å².

Preprocessing of these datasets was performed using RELION 4.0 (44). Final particles of τA-DNA and τB-DNA from each dataset were merged independently with a final refinement step in RELION 5 using blush regularization (45). CryoSPARC versions 3.3.2 to 4.4 were used throughout the data processing timeline. Supplementary Figure S4 and S5 contain detailed preprocessing methodology.

For the τB-DNA subcomplex analysis, particles picked by WARP from each dataset underwent 2-5 cycles of heterogeneous refinement in cryoSPARC , using a volume from an ab initio job based on specific 2D classes representative of the τB-DNA complex, along with 4-6 decoy volumes. Subsequently, particles were classified using cryoDRGN v2.3 using default parameters (46), and re-imported into RELION for further refinement, achieving resolutions between 3.09 Å and 3.75 Å. These particles, with optimized Euler angles, were combined into a dataset of 570,437 particles and processed in RELION 5 by using Blush regularization (45). Initial refinement yielded a 3 Å map, which was improved to an overall resolution of 2.46 Å after three rounds of CTF refinement and Bayesian polishing.

For the τA-DNA complex, each dataset was processed through a similar workflow. Initial particle picking was performed using WARP, followed by 2D classification. Subsequently, selected particles with τA-DNA features were imported and re-extracted in RELION at a 300 px box size for TOPAZ training and picking. The obtained set of TOPAZ-picked particles was extracted using a box size of 320 px, imported into cryoSPARC for classification through heterogeneous refinement and 2D classification to remove junk particles. These classified particle sets were then used for a second round of TOPAZ training and picking, followed by particle re-extraction and further heterogeneous refinement and 2D classification in CryoSPARC. The remaining particles from each dataset were merged for subsequent classification. A total of 739,929 particles were processed using an ab initio step to create a new τA-DNA map that was used in combination with other decoy volumes for two consecutive heterogeneous refinement steps. The τA-DNA map, containing 114,621 particles, was then imported into RELION5 for refinement using blush regularization (45). The final map showed an overall resolution of 3.65 Å after two rounds of CTF refinement and Bayesian polishing.

Despite of adding a range of transcription factors during sample preparation, the subsequent final maps of the τA-DNA and τB-DNA complexes did not exhibit any additional density that could be attributed to these added transcription factors.

### Model building, refinement and validation

Structural predictions for the τA and τB subcomplexes were generated using AlphaFold Multimer 2.3 (47). The τA subcomplex included the C-terminus of τ138 (aa 635 to 1060), τ131, τ95, and τ55 subunits. The τB subcomplex comprised the N-terminal region of τ138 (aa 1-668), τ91 and τ60. Highest-ranked predictions were converted from .pkl to .json files using a script from http://www.subtiwiki.uni-goettingen.de/v4/paeViewerDemo, enabling the utilization of AlphaFold prediction scores in ISOLDE (48).

Initial models for τA and τB subcomplexes were manually placed into their respective density maps and subjected to rigid-body fitting using ChimeraX (49). Using the .json files, models were refined in ISOLDE. This step was critical for accurately placing secondary structures and domains in regions of lower resolution (4.5 to 6 Å), particularly in the τA map, where *de novo* building was challenging. Iterative refinement was made to correct rotamer and Ramachandran outliers followed by real-space refinement using Servalcat (50) for both models. Validation of the refined models was conducted using MolProbity (51).

For the τB-DNA complex, a B-DNA model for the first 40 nucleotides of the downstream region of the ^His^tRNA gene was built using self-restraints in Coot (52). The upstream 45 base pairs for the τA-DNA complex were fitted into the DNA density, although the low resolution of this region might introduce some ambiguity regarding the A-box position and DNA orientation. Coot was also used to refine the protein-DNA interaction interface for both complexes. Protein-DNA interactions were analyzed using ChimeraX and DNAproDB (53,54). The DNA groove width was analyzed with Curves+ (55).

To obtain a structural model of the complete TFIIIC-DNA complex, the 85 bp ^His^tRNA DNA duplex was placed into the τB-DNA map with 40 bp (+37 to +76) fitting into the τB-DNA map. Subsequent refinement of the remaining 45 bps was performed in the τA-DNA map. Accurate positioning of the τA subcomplex relative to the DNA was achieved by integrating the orientations and distances derived from cryo-EM single particle mapping.

### Cryo-EM single particle mapping of TFIIIC

The independently refined RELION sets of coordinates and Euler angles for τA-DNA and τB-DNA particles from dataset 1 (see Supplementary Figure S3 – middle) provided the information necessary for distance pair and orientation mapping of both subcomplexes, at a single molecule level. For each micrograph, pairs of τA and τB particles belonging to an individual complex were identified by the nearest corresponding neighbor search. To avoid including the same particle in multiple pairs, this search was performed iteratively. The spatially closest particles were paired first and excluded for the following iterations in which the next closest pair was identified.

For each pair of τA and τB particles, the inter-particle distance was calculated from their XY-coordinates. The hypothetical distance distribution of unrelated particles was estimated by performing the same search on a theoretical dataset that maintained the numbers of particles per micrograph, but randomized their XY-coordinates.

The orientation of particles is described by using Euler angles. In RELION, the Euler angles α, β, γ (termed Rot, Tilt, and Psi in RELION) of each particle are determined by maximum likelihood regularized optimization using its respective 3D reference volume as a starting point. The refined Euler angles of τA and τB were used to derive their relative orientation.

To achieve this, the Python library scipy.spatial.transform.Rotation was employed. For each pair of particles, their absolute rotations (relative to their respective 3D reference) were converted from their Euler angle representation to scipy-rotation objects. The relative orientation of τA with respect to τB can be described with the rotation Rr. Rr was obtained by the combination of the inverted rotation of τA followed by the rotation of τB (RB * RA^-1^ = R_r_). This scipy-rotation object (R_r_) was then converted back to the Euler angles representation (Δαβγ_a→b_).

### Labeling of the ybbR-TFIIICΔtail complex with Cy5 for single-molecule fluorescence microscopy

The ybbR-TFIIICΔtail complex was labeled by Sfp-mediated coupling of CoA-Cy5 to the engineered ybbr-tag (56,57). We first tested various conditions to achieve optimal labeling efficiency. All reactions were performed in 20 µL volumes using a labeling buffer consisting of 50 mM HEPES pH 7.5, 100 mM NaCl, 20 mM MgCl_2_, and 5 mM DTT. The conditions varied primarily in enzyme concentration and incubation time, with ybbR-TFIIICΔ593 at 5 µM and CoA-Cy5 at 10 µM. Sfp enzyme concentrations were adjusted across trials (0.1 µM, 1 µM, and 2.5 µM), with incubation periods of 30, 60, and 90 min. Additionally, the impact of temperature was evaluated by conducting one set of reactions at 37°C, while the rest was kept at 25°C. Each condition was assessed by splitting the reaction mixtures: half were analyzed by SDS-PAGE and visualized on a Typhoon scanner using the Cy5 channel, and the other half were evaluated for functionality using an electrophoretic mobility shift assay.

The labeling was scaled up to a 500 µL reaction volume with following final reaction conditions: 5 µM ybbR-TFIIICΔtail complex, 2.5 µM Sfp enzyme, 10 µM CoA-Cy5 dye incubated for 30 min at 25°C. Excess dye was removed post-reaction through buffer exchange with a PD-10 desalting column. The eluates were then analyzed by SDS-PAGE to ensure minimal dye contamination, with the cleanest fractions pooled and subjected to a second purification step using another PD-10 column. Absorbance of the purified fractions was measured at 280 nm and 646 nm to assess protein concentration and labeling efficiency.

### Preparation of biotinylated and fluorescently labelled DNA for single-molecule fluorescence microscopy

To prepare DNA constructs A/B, mA/B, mA/mB and A-only for single-molecule fluorescence microscopy (SMFM), adapter sequences were introduced at both termini. The 5’ overhang was used to hybridize a Cy3-labeled DNA oligo (p0074-Cy3 oligo). The 3’ overhang was designed to hybridize a biotin oligo (p0141-p0109-biotin) for immobilization of the tRNA gene sequence to the glass slide surface (see Table S2 for sequence details).

The DNA template was generated by autosticky PCR as described previously (58). In short, the DNA template with single-stranded overhangs was generated with primers containing an abasic site. Following electrophoresis, the DNA templates were gel-purified and subjected to buffer exchange into SMFM DNA buffer (10 mM Tris-HCl pH 7.5, 20 mM KCl) using an Amicon Ultra-0.5 Centrifugal Filter with a 3 kDa MWCO, concentrating to a final volume of around 1 µM. 100 nM DNA template was incubated with 120 nM p0074-Cy3 and 100 nM pre-annealed p0141-p109-biotin oligos in 10 mM Tris-HCl pH 7.5, 20 mM KCl for 5 min at 68°C followed by the slow cool down.

### Single-molecule experiments to detect TFIIIC binding dynamics

To detect binding dynamics of TFIIIC-Cy5, 50 pM of Cy3-DNA-biotin was immobilized on biotin-polyethylene-glycol (PEG) functionalized glass slides, priorly coated with NeutrAvidin for 10 min at room temperature (59). After 5 min immobilization, the unbound DNA molecules were washed away with imaging buffer (SMFM reaction buffer containing additionally 0.25 % Biolipidure 203, 0.25 % Biolipidure 206, an oxygen scavenger system: 2.5 mM protocatechuic acid, 100 nM protocatechuate dioxygenase, and a mix of triplet state quenchers: 1 mM 4-citrobenzyl alcohol (NBA), 1 mM cyclooctatetraene (COT) and 1 mM Trolox). The experiment was initiated by delivering 20 nM TFIIIC-Cy5 and 1 µM competitor DNA (as poly(deoxyinosinic-deoxycytidylic) acid sodium salt, Merck) in the imaging buffer ∼15 sec after the start of the single-molecule imaging.

### Instrumentation for single-molecule fluorescence imaging and data analysis

Single-molecule experiments were performed at 21°C on a custom-built objective-based (CFI SR HP Apochromat TIRF 100XC Oil) total internal reflection fluorescence (TIRF) microscope (built by Cairn Research: https://www.cairn-research.co.uk/). The TIRF microscope was equipped with an iLAS system (Cairn Research) and Prime95B sCMOS cameras (Teledyne Photometrics). The MetaMorph software package (Molecular Devices) was used for data acquisition. For alternative laser excitation (ALEX) experiments, the immobilized sample was excited with a diode-based (OBIS) 532 nm laser at 0.73 kW cm^-2^ output intensity (200 ms exposure time) in every odd frame and with a diode-based 638 nm laser (Omicron LuxX) at 0.24 kW cm^-2^ output intensity (200 ms exposure time) in every even frame. For non-ALEX experiments, which were used for all quantitative evaluation, we used the 532 nm laser at 0.86 kW cm^-2^ output intensity (200 ms exposure time).

To extract and process single-molecule traces from acquired movies, we used the SPARTAN software package (v.3.7.0) (60). To analyze the FRET efficiency distributions during TFIIIC-Cy5 binding events, we first used SPARTAN to select traces with a single Cy3 photobleaching step and to perform background, spectral crosstalk and relative donor/acceptor scaling corrections. For further processing, the traces were exported to the tMAVEN software (61). The evaluation windows were selected to include only the TFIIIC-bound regions. To model the number of FRET states we used hidden Markov modeling (global vbConsensus + Model Selection) in tMAVEN and used MATLAB to fit FRET efficiencies distributions to Gaussian functions using the maximum likelihood function in MATLAB. For analysis of the TFIIIC bound lifetimes, traces were exported from SPARTAN and then analyzed by MATLAB (version R2019) (62,63). The TFIIIC-Cy5 bound dwells were fitted to a single or double-exponential function to extract the lifetimes. The error bounds reported in the text represent the standard deviation from at least two independent experiments.

### Protein and DNA sequence analysis

The sequence identity between *S. cerevisiae* τ138 (Uniprot ID: P34111) and *H. sapiens* TFIIIC220 (Uniprot ID: Q12789) was calculated via the UniProt ’Align’ tool (64). The conservation of the amino acids that contribute to base-specific readout of the B-box was inferred from a multiple sequence alignment (MSA) generated by us in an earlier conducted study (32). For the analysis of the ETC loci, the sequence identifiers (ETC-ETC8, ZOD1) were taken from Moqtaderi et al. (18). The corresponding sequences were fetched from the Saccharomyces genome database (SGD) (65) and aligned with the Muscle software (66). The region corresponding to the ETC B-box was selected based on the analysis by Moqtaderi et al. (18). For comparison with the canonical B-box in tRNA genes, the *S. cerevisiae* tRNA genes (dataset: *Saccharomyces cerevisiae* S288c) were downloaded from the GtRNAdb 2.0 database (Release 21, Jun 2023) (67). The sequences were aligned with *cmalign* program that is implemented into the Inferal package (68) using the tRNA covariance model RF00005 as input. This covariance model was retrieved from the Rfam database (69). Sequence logos of the canonical tRNA- and the ETC B-boxes were then generated via WebLogo (70).

## Results

### Cryo-EM structures of DNA-bound τA and τB subcomplexes

*Saccharomyces cerevisiae* TFIIIC comprises two evolutionarily conserved modules: τA contains the C-terminal region of τ138 and subunits τ131, τ95, and τ55; and τB contains the N-terminal region of τ138 and subunits τ91, and τ60 (32). All TFIIIC subunits were heterologously co-expressed in insect cells as full-length proteins, except of subunit τ95 that was C-terminally truncated at residue 593 to enhance τA interaction with the A-box by removing an acidic tail that decreases DNA binding (9). To verify the structural integrity of the purified TFIIIC complex, referred to as TFIIICΔtail, we performed mass photometry (MP). This analysis determined a molecular mass distribution with a main peak at 547 ± 35 kDa, closely matching the theoretical molecular weight of 520 kDa (Supplementary Figure S1B). Furthermore, a filter binding assay confirmed the high affinity of TFIIICΔtail for the 85 bp ^His^tRNA gene, with an apparent dissociation constant (K_D_) of 30 nM (Supplementary Figure S1C).

Upon determining the optimal conditions for cryo-EM, an initial dataset, referred to as dataset 1, comprising 19,047 micrographs, was collected. Early stages of data processing identified three distinct sets of particles: τB-DNA dimers, τB-DNA monomers, and τA-DNA monomers (Supplementary Figure S3). The τB-DNA monomer map achieved the highest resolution of 3.21 Å, while the τB-DNA dimer displayed significant anisotropy (Supplementary Figure S3). The τA-DNA monomer map had lower resolution due to the dynamic interaction of τA with DNA and fewer particles in a consistent binding state (Figure 2A). To improve resolution, four additional datasets (datasets 2-5) were collected (Supplementary Figures S4 and S5) that in addition to TFIIIC contained additional factors namely TFIIIB, TBP-Brf1 and Fpt1 although none of these factors could be detected in the final cryo-EM maps (Supplementary Table S1).

Processing all datasets yielded cryo-EM maps of the *S. cerevisiae* TFIIIC-type II promoter complex capturing both τA and τB subcomplexes in DNA-bound states (Figure 1B). This contrasts with the previously characterized human TFIIIC, which only showed τB engaged with DNA (32). Separate cryo-EM reconstructions of the τB and τA subcomplexes were achieved at 2.46 Å and 3.65 Å resolution, respectively (Supplementary Figures S4 and S5). The quality of the obtained cryo-EM maps enabled building and refinement of atomic models for both subcomplexes (Figure 1C, Supplementary Figure S6; Table S1a and S1b).

**Figure 1.**
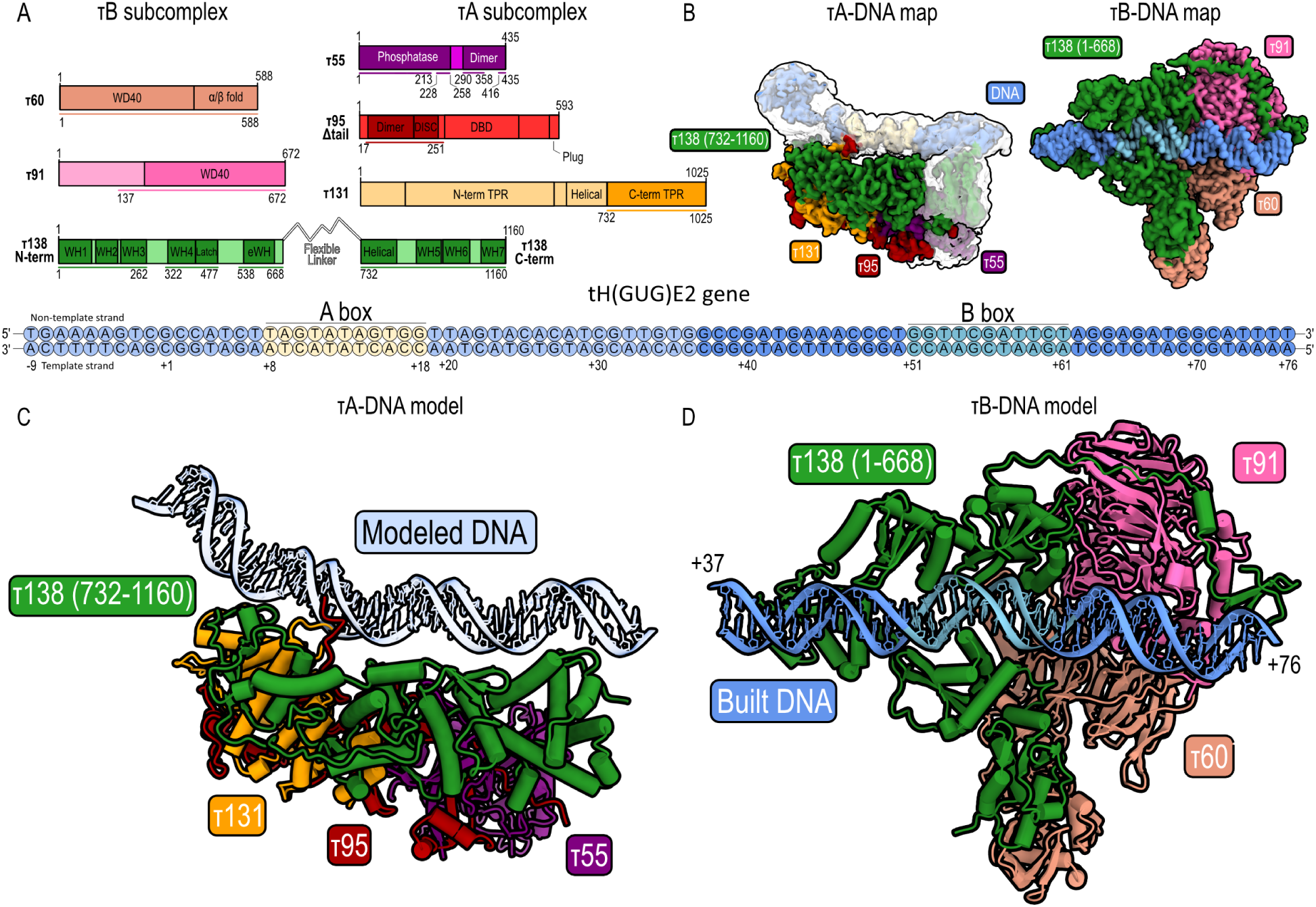
Cryo-EM structures of TFIIIC bound to tRNA gene. (**A**) Domain diagram of yeast TFIIIC subunits divided into τA and τB subcomplexes, where colored bars represent built regions (left). Key domains include: DBD - DNA binding domain, TPR - tetratricopeptide domain, WD40 - WD40 repeat domain, WH - winged-helix domain, eWH - extended winged-helix domain. The ^His^tRNA gene, used in the cryo-EM study (right), is depicted with fully colored circles highlighting the DNA-interacting τB subcomplex (built DNA), whereas the rest of the DNA is represented by transparent colored circles (modeled DNA). (**B**) Cryo-EM maps of τA-DNA and DNA-bound τB subcomplex. Superposition of local resolution τA-DNA map (transparent) on the corresponding sharpened τA-DNA map to reveal densities sufficient for modeling 38 bp of DNA, helical domain (τ138) and phosphatase domain (τ55) (**C**) Cryo-EM structure of τA-DNA complex. (D) Cryo-EM structure of τB-DNA complex.

**Figure 2.**
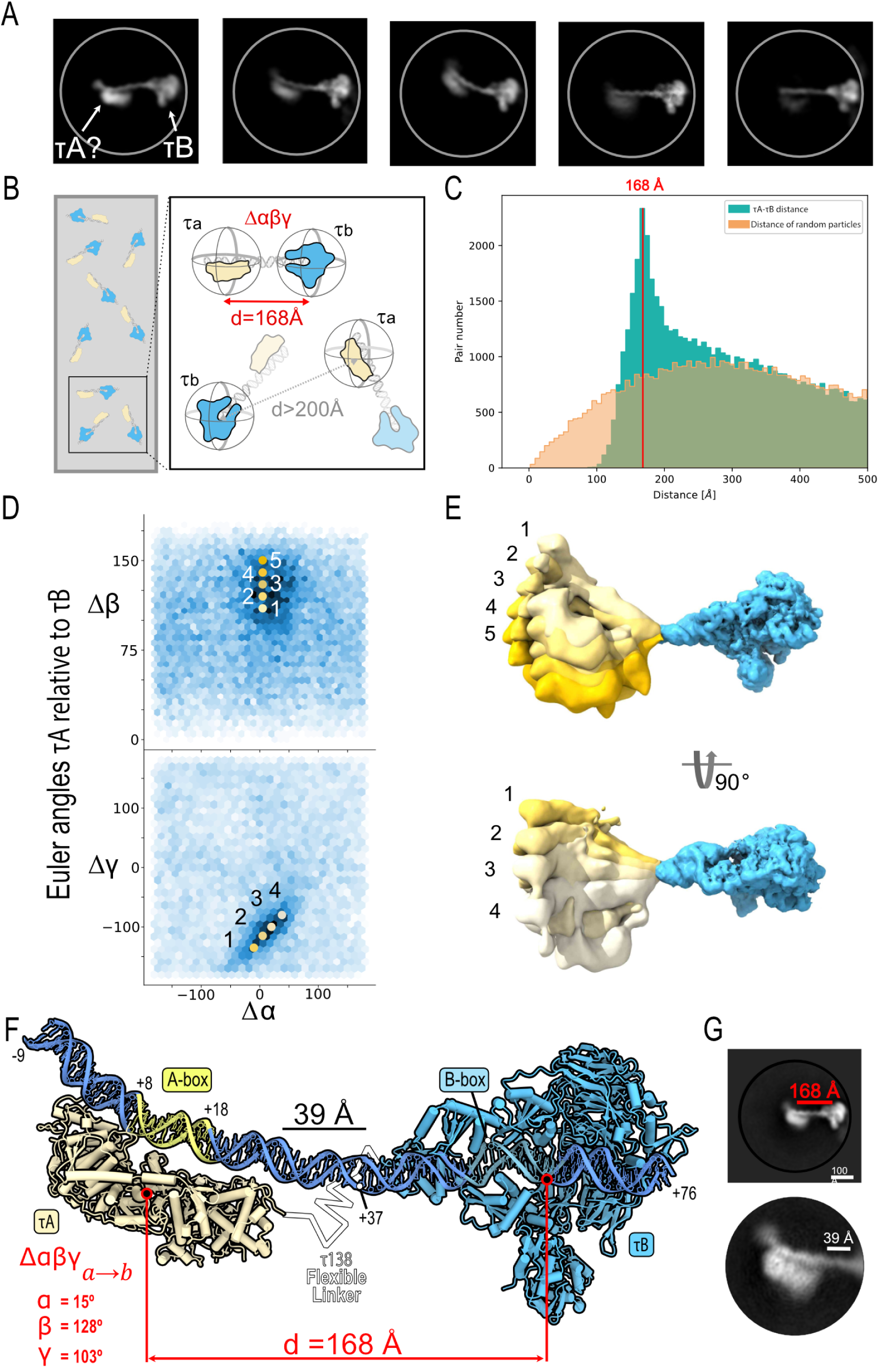
Cryo-EM single particle mapping reveals TFIIIC-promoter architecture. **A)** 2D classes from dataset 1, recentered along the DNA downstream of the τB-bound B-box. The class averages reveal cryo-EM densities of a continuous stretch of DNA bridging the τB-DNA subcomplex and the putatively assigned τA-DNA subcomplex. **B)** Independently refined particle coordinates and Euler angles of τA and τB particles were used to map the distances and orientations of τA and τB in complex with ^His^tRNA. Distances were mapped by a nearest neighbor search, while the orientation of τA relative to τB (Δαβγ) was calculated for the identified complexes. **C)** Mapped distance distribution between τA-DNA and τB-DNA (green) is compared to distance distribution of simulated particles with random coordinates (orange). A distinct peak suggests an average distance of 168 Å between τB and τA when bound to DNA (marked in red). **D)** Angular space of τA and τB described by relative Euler angles Δα, Δβ, Δγ. All τA-τB pairs with distance <200 Å were plotted. An enriched population around Δα ≈ 15°, Δβ ≈ 128°, and Δγ ≈ -103° was obtained. Representative values describing the spread of the population were selected (yellow dots: upper Δαβγ (5, 110, -115); (5, 120, - 115); (5, 130, -115); (5, 140, -115); (5, 150, -115) and lower Δαβγ (-10, 130, -135); (5, 130, -115); (20, 130, -100); (38, 130, -80)) and applied to the cryo-EM volume maps in panel e. **E)** Range of motion observed in the single-particle level mapping. Conformations correspond to the indicated positions in angular space. Indicated orientations were applied to the τA volume (yellow). τA and τB (blue) were spaced by 168 Å. For both subcomplexes the cryo-EM volume maps of dataset 1 refinement were used. **F)** Integrated structural model of full TFIIIC complex bound to A- and B-Box on basis of the cryo-EM structures τA and τB subcomplexes, 85bp ^His^tRNA dsDNA model, and cryo-EM single-particle mapping of distances and orientations. The experimentally determined DNA structure is opaque, whereas the modeled DNA is transparent. The A-Box colored in pale yellow and B-Box colored in teal. **G)** The 2D class average from reextracted particles of pairs with a distance < 200 Å shows the full TFIIIC-DNA complex (500 Å mask diameter). They illustrate the TFIIIC complex in association with the tRNA gene, displaying variability in the clarity of the τA subcomplex while maintaining a consistently well-defined τB-DNA subcomplex.

### Structural architecture of the TFIIIC-tRNA promoter complex

Next, we asked whether TFIIIC can adopt a fully engaged state in the context of a ^His^tRNA gene, simultaneously recognizing its A- and B-box. To visualize the entire TFIIIC-DNA complex, we initially performed 2D classifications of particles from dataset 1, which yielded the reconstruction of the τB-DNA subcomplex at 3.2 Å resolution (Supplementary Figure S3 – middle). We re-centered the particles along the DNA downstream of the τB-bound B-box and extracted them using an extended box size (493 Å). The 2D class averages revealed a continuous stretch of DNA protruding from the τB-DNA subcomplex, with densities, opposite τB, which may correspond to the DNA-bound τA subcomplex (Figure 2A). However, structural heterogeneity prevented us from obtaining 3D reconstructions of the entire complex.

To accurately determine the 3D spatial arrangement of the TFIIIC-DNA complex, we employed cryo-EM single-particle mapping (Figure 2B, Supplementary Figure S7). This approach utilizes independently refined subcomplexes and has recently enabled us to characterize, at a single-molecule level, the structural heterogeneity of similarly sized complexes including human TFIIIC and the hexasome-INO80 complex (32,71). A nearest-neighbor distance search of the DNA-bound τA and τB subcomplexes from dataset 1 identified a distinct population of particle pairs with a peak at a distance of 168 Å (Figure 2C). This mapping allowed us to determine the relative orientations of τA and τB subcomplexes in the context of a single ^His^tRNA gene. By utilizing independently refined Euler angles of each subcomplex, our analysis revealed a distinct range of DNA-bound τA orientations relative to τB (Figure 2D,E, Supplementary Figure S7). Integrated structural modeling based on the determined cryo-EM structures of DNA-bound τA and τB, the 85bp ^His^tRNA sequence as well as the mapped distance and orientation distribution visualizes a fully engaged state of TFIIIC (Figure 2F). Re-extraction of identified particle pairs combined with further 2D classification validated the obtained model (Figure 2G), while the observed structural heterogeneity of the complex can be explained by the flexible bending of the linker DNA (Supplementary Movie 1). Notably, τA is mapped at a defined distance and orientation range that suggests specific A-box recognition (Figure 2F).

### B-box recognition by the yeast TFIIIC τB subcomplex

The yeast TFIIIC τB subcomplex bound to a type II promoter described in this study resembles human and yeast TFIIIC τB bound to type II and type I promoters, respectively (32,33). Specifically, the yeast τB subcomplex includes the N-terminal region of τ138 (1-668 aa), τ91 and τ60 (Figure 1). The N-terminus of τ138 comprises several distinct domains: WH1 (aa 1-97), WH2 (aa 108-172), WH3 (aa 185-262), WH4 (aa 334-416), and eWH (aa 550-640). Like for the human τB subcomplex, we show that the first domain of τ138 is a WH domain and not a High Mobility Group (HMG) domain in contrast to earlier analyses (27,33,35).

The high resolution of 2.46 Å for the τB-DNA subcomplex facilitated the accurate building of 40 bp (+37 to +76) from the 3’ end of the yeast ^His^tRNA gene (Figure 1C). Unlike in the human counterpart, all WH domains in yeast interact with the downstream region of the ^His^tRNA gene through base-specific contacts (Figure 3A). Specifically, WH2 interacts with the B-box motif through residue R139 contacting C55 on the non-template strand as well as G55 and C56 on the template strand; residue S140 also interacts with C56 on the template strand and residue H162 interacts with G51 on the non-template strand (Figure 3B). In WH3, residue K223 forms base-specific hydrogen bonds with G47 on the template strand, a region upstream the B-box (Figure 3A and B). This interaction is notable as it is the only one among the WH domains in the N-terminus of τ138 that extends outside the B-box. This observation aligns well with findings from studies on other tRNA genes, such as ^Tyr^tRNA or SUP4, where a mutation in a nucleotide upstream of the B-box (G45 to A45) led to a five-fold increase in the equilibrium constant for TFIIIC binding (11).

**Figure 3.**
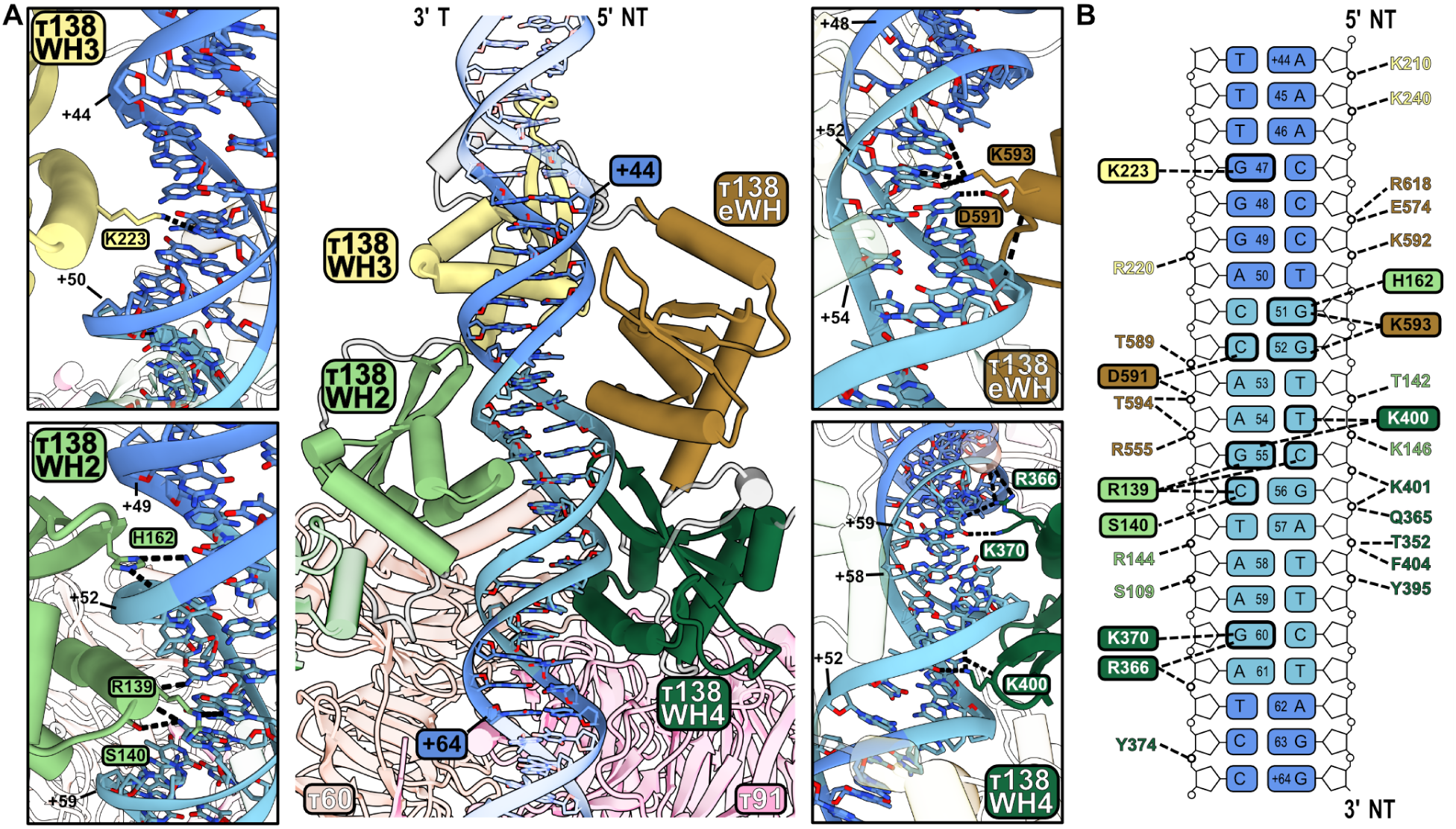
Interaction of the N-terminus region of τ138 subcomplex with the B-box. (**A**) Cryo-EM structure of the τB-DNA subcomplex, highlighting in full color the τ138 domains that interact with DNA. The B-box motif is shown in light blue. The interactions between WH2, WH3, WH4, and eWH domains with DNA are shown in detailed views. (**B**) Schematic of the interactions, including all hydrogen bonds, between the τ138 subunit and the tRNA gene. Hydrogen bonds are represented by black dashed lines. Amino acids that make contact with DNA bases are presented in colored boxes.

Further, site-directed mutagenesis in tRNA genes from *C. elegans* (72) and *S. cerevisiae* (73) demonstrated that transcriptional activity changes when nucleotides from +44 to +47 were altered. Our results demonstrate the importance of nucleotide positions outside the B-box, highlighting their crucial role in modulating TFIIIC binding affinity and transcriptional activity. Prior dimethyl sulfate protection experiments identified several guanine bases in the ^Glu^tRNA_3_ B-box as crucial for the formation of the yeast τB-DNA complex (74). These guanine bases (G53 on the non-template strand and G56, G61, and G62 on the template strand) correspond to G52 on the non-template strand and G55, G60, and A61 on the template strand of the ^His^tRNA gene. For the WH4 domain, residues R366 and K370 interact with G60 on the template strand, and K400 interacts with T54 on the non-template strand and G55 on the template strand (Figure 3A and B). While eWH’s residue D591 interacts with C52 on the template strand and K593 interacts with G51 and G52 on the non-template strand (Figure 3A and B).

For comparison, the cryo-EM structures of the τB subcomplexes bound to DNA from yeast and human were superimposed using the B-box sequence region (Supplementary Figure S8A). Despite only 18.7% sequence identity between the yeast and human τ138 subunits, the structural configuration surrounding the B-box, encompassing all WH and eWH domains, is strikingly conserved. Furthermore, this superimposition reveals that the WH2 and WH4 domains in human and yeast diverge the least in terms of their spatial positioning, as opposed to WH3 and eWH. This observation aligns with the roles of WH2 and WH4 as part of the τB scaffold for initial DNA recognition, anchored to the τB core, which also comprises the τ60 and τ91 subunits in yeast (32). A comparison of the key residues in yeast and human involved in base-specific interaction with the downstream region of tRNA is also analyzed (Supplementary Figure S8B). This conservation is also evident in the altered conformation of the B-box DNA in both yeast and human structures (Supplementary Figures S8C). In these structures, the minor groove at position C55/C69 (yeast/human) is significantly widened, while it is notably narrowed near T59/T73. Additionally, several of the B-box contacting residues (highlighted in pink) are either identical or highly conserved across a wide range of eukaryotes, including metazoans, fungi, amoebozoa, plants, and discobians (32).

To gain further insights into the role of TFIIIC in binding type I and type II promoters, a comparison of τB-DNA complex structures was performed. The flexibility of the WH3 and eWH domains becomes apparent when examining TFIIIC in the context of the type I promoter in the presence of TFIIIA (Supplementary Figure S9A). Including TFIIIA alters the interactions of the WH domains and eWH with DNA. On type II promoters, all WH domains and eWH engage with DNA (Supplementary Figure S9B). However, TFIIIA’s presence strongly displaces WH3 and eWH and to a smaller extent WH2 (Supplementary Figure S9C – arrows color-coded to match the domains indicate their movement) leaving only WH4 seemingly unaffected by this additional factor. Overall, this comparison shows the pivotal role of TFIIIA in modulating the conformation of TFIIIC, particularly influencing the arrangement of the WH domains and eWH in the τB-DNA complex, and highlights the flexibility of these domains in response to different transcriptional requirements in Pol III genes.

### The τ60/τ91 heterodimer contacts DNA bases outside the B-box

The yeast τB core comprises the τ60/τ91 heterodimer and closely resembles the human τB core with subunits TFIIIC110 and TFIIIC90 forming the heterodimer (32). A truncated version of the heterodimer τ60/Δτ91 (aa 159-672) has been studied using X-ray crystallography that showed no DNA-binding activity (34). In our τB-DNA structure, subunit τ91 is binding to the downstream region of the B-box with residues K591 and N628 making base-specific contacts (Supplementary Figure S10A). Although these amino acids are also present in the τ60/Δτ91 structure, the absence of residues 137 to 158 in construct Δτ91, especially K153 and R155 that interact with the phosphate backbone of the DNA might be important for stabilizing this interaction. In addition, the N-terminal region of τ138, which is part of the τB subcomplex, might also help τ91 binding to DNA. In human TFIIIC, the positively charged surface of the TFIIIC110 subunit contributes to the interaction with DNA (32). A study that first cloned and characterized τ91 found that this subunit cooperates with τ138 for DNA binding (15). Additionally, τ60 in yeast contributes to DNA binding by engaging with the phosphate backbone of the B-box motif through residues R11 and S362 (Supplementary Figure S10B).

The interactions of subunit τ91 with DNA-bases downstream of the B-box is conserved between 9 ETC loci where TFIIIC presumably functions as genome organizer (18) (Supplementary Figure S10C). TFIIIC’s role as genome organizer has been also hypothesized to be linked with the ability of human τB to dimerize (32). Yeast τB also dimerizes under conditions used for cryo-EM grids preparations. Two types of dimers were identified that were termed “thumb-knuckle” and “knuckle-knuckle” to distinguish their interface interactions were identified (Supplementary Figure S11A). Mass photometry experiments assessed TFIIIC dimerization *in vitro* with and without DNA (85 and 120 bp ^His^tRNA DNA) and revealed TFIIIC dimers at 50 nM protein concentration indicating that TFIIIC dimerization is concentration-dependent (Supplementary Figure S11B).

### A-box recognition by the yeast TFIIIC τA subcomplex

The domain architecture of yeast τA bound to DNA is depicted in Figure 4A. It includes the τ138 C-terminal region, which is integral to the formation of the τA subcomplex due to its interactions with other subunits (Figure 4B - top). Overall, the structure of yeast τA is similar to human τA with notable variations (32). In yeast τA, subunit τ138 contributes with one helical domain and only three WH domains (WH5 – WH7), whereas in human τA, the corresponding subunit TFIIIC220 contributes one homeobox-like domain and four WH domains (WH5 – WH8) (Figure 4B). Additionally, only the C-terminal TPR domain of τ131 is resolved in yeast, unlike in human TFIIIC102, where part of the N-terminal TPR domain is also resolved. This difference can be attributed to the flexibility of τ131 and its role in ’fishing’ for TFIIIB once τA is bound to DNA. Yeast subunit τ55 and its human counterpart TFIIIC35 dimerize with subunit τ95 and corresponding human subunit TFIIIC63 through a conserved triple β-barrel domain. Likewise, yeast τ95 and human TFIIIC63 both contain a disc domain showing the structural similarity between species. In contrast, yeast τ55 additionally contains a phosphatase domain (36) that is absent in TFIIIC35.

**Figure 4.**
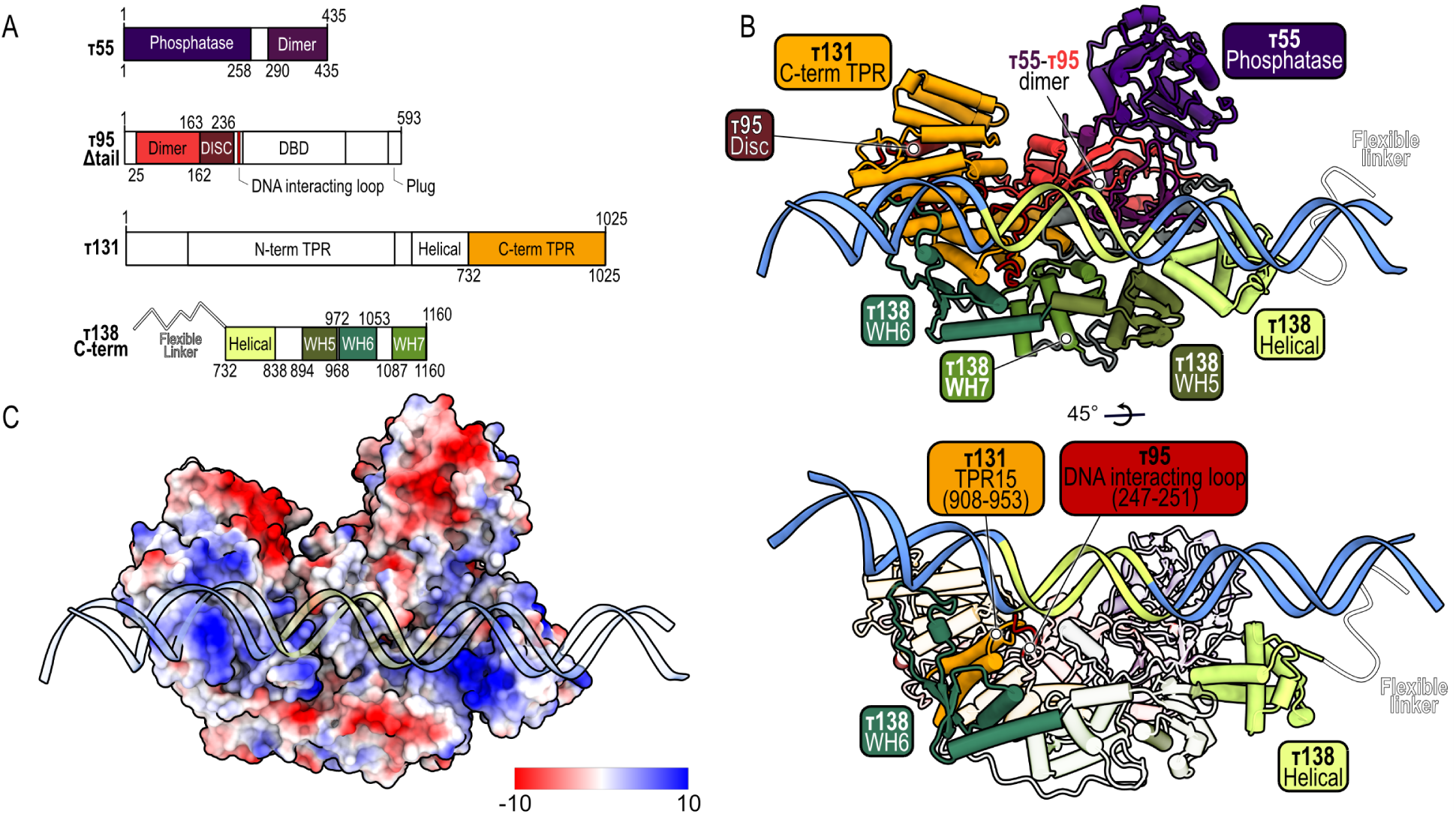
Different τA subunits interact with DNA. (**A**) Domain architecture color-coded to match the cryo-EM model in panel B, with only built domains colored and the rest shown in white. (**B**) Cryo-EM model of τA-DNA complex, detailing domain composition with the flexible linker indicated by a white line (top). Cryo-EM model showing only the domains involved in DNA interaction in full color (bottom). (**C**) Electrostatic potential surface of τA subcomplex, with DNA shown in a cartoon and transparent style for clarity.

In yeast τA, four domains interact with the ^His^tRNA DNA: the helical domain (aa 732 to 838) and WH6 (aa 972 to 1053) in τ138, TPR15 in τ131 (aa 908 to 953), and the DNA-interacting loop (aa 247-251) in τ95 (Figure 4B – bottom). Although several τA domains are involved, each domain establishes only a relatively small interface, mostly via electrostatic interactions with the DNA phosphate backbone (Figure 4C). This binding mode may account for the weak DNA binding affinity of τA, potentially becoming stabilized by additional factors like TFIIIB (7). However, even in the absence of TFIIIB (dataset 1), TFIIIC specifically recognizes the A-box DNA motif: cryo-EM single particle mapping unambiguously demonstrated that τA adopts a specific orientation on a ^His^tRNA DNA directly at the A-box motif (Figure 2). Notably, when we probed a slightly longer ^His^tRNA DNA with a putative second A-box (dataset 2 - 5), we noticed some re-distribution of τA to this second motif. While the relatively low resolution of DNA in the τA subcomplex did not allow us to assign a more precise register of DNA sequence recognition, we observed a pronounced overall DNA curvature suggesting DNA shape recognition. Notably, the loop insertion of τ95 (aa 247-251) and τ131 (aa 908 to 953) into the minor groove of the A-Box might contribute to a combined DNA sequence and shape readout (Figure 4B), but based on our cryo-EM analysis we cannot exclude contributions by other, more dynamic interactions.

In the τA-type II promoter complex, the ^His^tRNA gene spans the τA subcomplex, crossing the helical domain at its most downstream point and extending directly to the WH6 domain of τ138 at its most upstream region. The A-box is in close contact with the τA subcomplex (Supplementary Figure S12A - left). In contrast, in the type I promoter (5S rRNA gene) with TFIIIA, the DNA adopts a perpendicular orientation relative to that observed in type II Pol III genes. Here, the main DNA-protein interactions involve the C-terminal and N-terminal TPR domain repeats of τ131, with no direct interaction with the A-box (Supplementary Figure S12A - right). Thus, TFIIIA significantly changes the way τA interacts with its target DNA (Supplementary Figure S12A). Additionally, among all the regions and domains involved in DNA binding in type I and II promoters, only the interaction involving TPR15 of subunit τ131 is common to both (Supplementary Figure S12B).

### Single-molecule fluorescence microscopy shows dynamic interactions of TFIIIC with DNA in real time

Upon analyzing the 2D classes of the τA-DNA complex in detail, a fuzzy region can be observed at the most upstream region of the DNA, appearing in different positions relative to the τA core (Figure 5A). These 2D classes also reveal dynamic contacts of τA and the DNA at the opposite site of the fuzzy region. This fuzzy region likely corresponds to the putative τ95 DBD, which is absent in our τA-DNA structure. In *S. pombe*, it has been hypothesized that the τ95 DBD requires bent DNA to avoid steric clashes and optimize interaction with double-stranded DNA (12). Consistent with this hypothesis, our τA-DNA cryo-EM structure reveals bent DNA in the upstream region (Figures 1C and 4B), as well as in the 2D classes (Figure 2C and Figure 5A).

**Figure 5.**
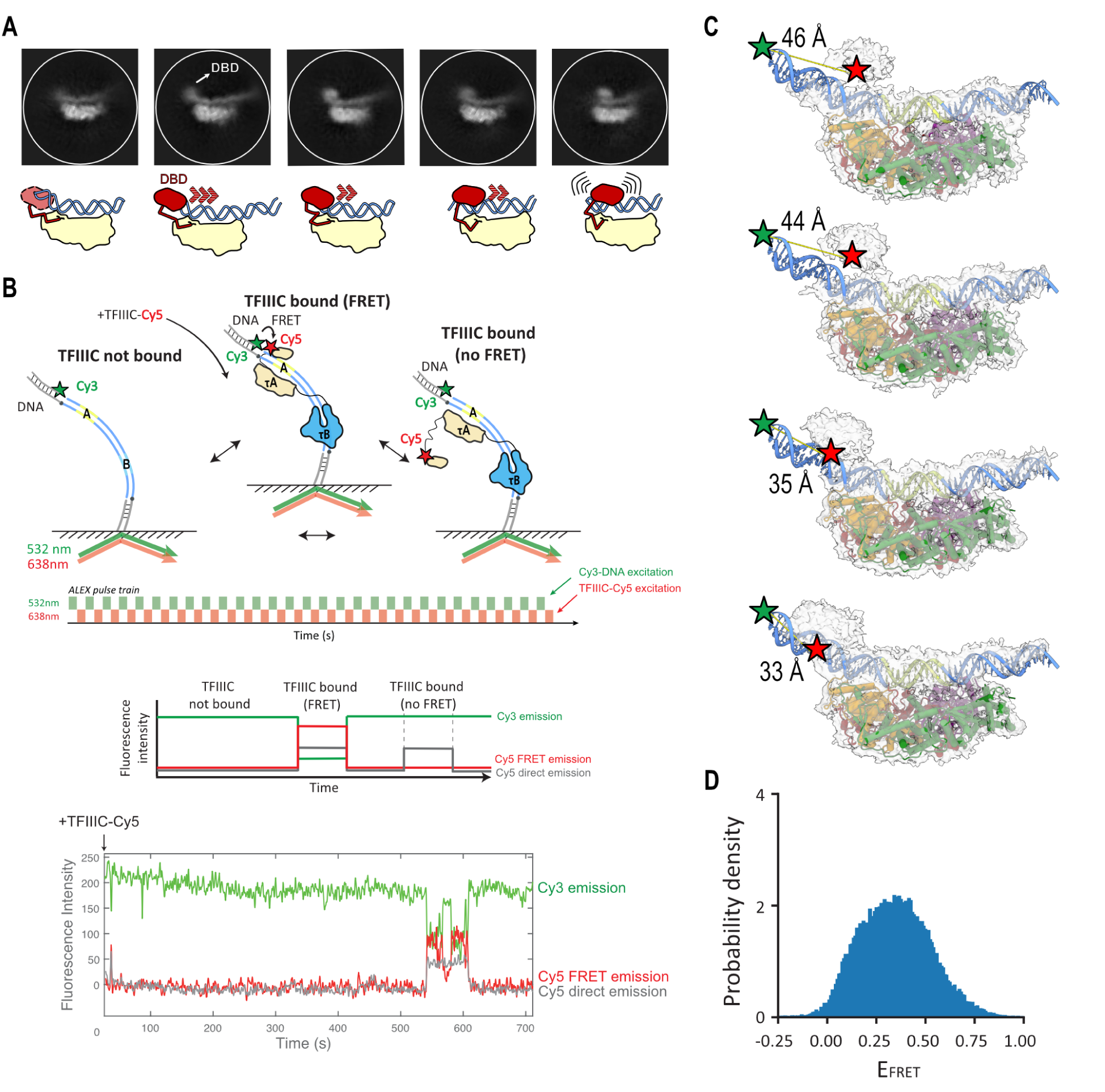
DBD/τA conformational dynamics. **(A)** Top: Selected 2D classes from dataset 1 showing τA bound to DNA with a fuzzy region, presumably the DBD-τ95, moving along the DNA. Bottom: A schematic representation of the selected 2D classes, illustrating the potential movement of the DBD (red) relative to the τA core (yellow). **(B)** Upper part: experimental setup to detect TFIIIC-Cy5 binding to Cy3-labeled DNA template. Central part: schematic representation of a single-molecule trace. Bottom part: experimental single-molecule trace showing TFIIIC-Cy5 binding to A/B DNA-Cy3 (TFIIIC-Cy5 bound in FRET distance – red, all TFIIIC-Cy5 bound – grey). (**C)** Cryo-EM maps generated by cryoDRGN along PC1. Approximate distances between the Cy3 dye (green star) on the DNA and Cy5 (red star) on the density corresponding to the DBD are indicated. **(D)** FRET efficiency distribution of TFIIIC-Cy5 bound to A/B DNA, 278 dwells analyzed, number of datasets = 2.

Given the possible dynamic nature of the τA-DNA interactions identified through our structural analyses, we sought to directly track the binding dynamics of TFIIIC with DNA in real-time using single-molecule fluorescence microscopy. To this end, we immobilized a Cy3-tagged DNA template containing both A-box and B-box (A/B oligonucleotide) to a functionalized glass surface for single-molecule imaging (Figure 5B). About 15 seconds after the start of the experiments, we delivered 20 nM TFIIIC, which was Cy5-labelled at the τ95 DBD domain using ybbr-mediated dye conjugation (ybbr-peptide inserted between residue 286 and 287, see methods). Informed by our structural data (Figure 5C), the Cy3 donor dye on the DNA (labeled at position -14) and the Cy5 acceptor dye on τ95 DBD have an approximate distance of ∼ 33-46 Å and thus, fully DNA-engaged TFIIIC would result in detection of a high Cy3-Cy5 FRET signal upon 532 nm laser illumination. In order to also detect TFIIIC binding events, in which the dynamic τ95 DBD domain is positioned differently or is not even bound to DNA, we also directly excited the bound Cy5-TFIIIC molecules by additional 638 nm laser illumination using alternative laser excitation (switching 532 nm and 638 nm lasers excitation every 200 ms, Figure 5B).

The majority of the TFIIIC binding events showed FRET (Figure 5B, Supplementary Fig. 13A), indicating that if TFIIIC binds, it does so by binding to its specific binding sites, simultaneously (or at least within 200 ms) engaging with both A-box and B-box rather than sequentially binding to the B-box followed by binding to the A-box, as suggested in the human TFIIIC model (32). Interestingly, during a bound TFIIIC event, the Cy3-Cy5 FRET efficiency transitions between various states (Figure 5B, Supplementary Fig. 13A) resulting in a broad FRET efficiency distribution (Figure 5D). This FRET efficiency reports on the distance between the Cy3-label placed at the 5’end of the DNA template and the Cy5-label on the τ95 DBD of TFIIIC. Our smFRET data do not allow us to unambiguously assign whether these various states correspond to i) different DBD-bound conformations (Supplementary Figure 13B; possibility 1 and 2), ii) conformations with DBD-unbound but rest of the τA module DNA-bound (Supplementary Figure 13B; possibility 3), iii) a completely dissociated τA module conformation (Supplementary Figure 13B; possibility 4) or iv) other possible states. However, the cryo-EM data suggest that the various FRET states we observe at the single-molecule level represent various τ95 DBD-bound conformations with the τ95 DBD domain positioned differently with respect to the rest of the τA module (Figure 5C). While hidden Markov modeling of the various FRET states predicts two states as the most probable description of the data (Supplementary Figure 13C), there are likely more than two states present in solution as suggested by the cryo-EM structural analysis (Figure 5C), but with inter-dye distance differences too small to be assigned to more states.

We next quantified the TFIIIC-bound lifetimes and obtained a median TFIIIC-bound lifetime of 2+/- 1 seconds (Figure 6A,B). However, we see both short and long-lived binding events that show FRET (Figure 6A, top left panel), suggesting at least two binding modes in which both A-box and B-box are engaged. Fitting the bound dwell times to a double-exponential function, we obtain

**Figure 6.**
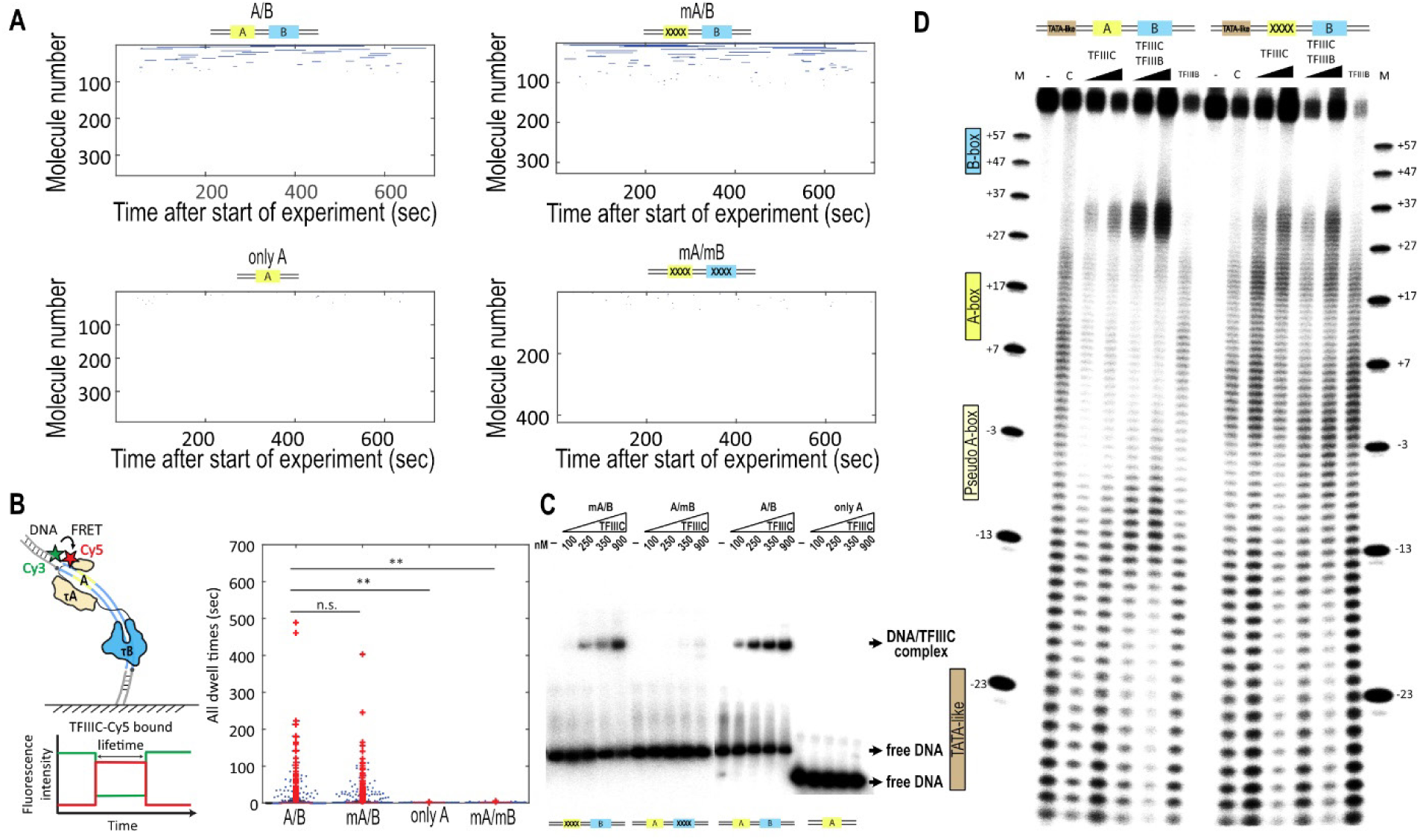
TFIIIC binding dynamics. **(A)** Rastergram of 20 nM TFIIIC-Cy5 binding to different DNA templates: for A/B DNA the number of molecules analyzed is n= 355; for mA/B DNA n = 327; for A-only DNA n = 392; for mA/mB DNA n = 423. (**B)** Left panel, schematic representation of analyzed TFIIIC-Cy5 bound lifetime. Right panel, beeswarm and boxplot plot of TFIIIC-Cy5 bound dwell times to A/B, mA/B, A-only, mA/mB. Number of evaluated molecules and number of analyzed dwells: A/B DNA - 771 and 188 (number of data sets = 2); mA/B DNA - 644 and 347 (number of data sets = 2); A-only - 392 and 45; mA/mB - 423 and 59. T-test: not significant (n.s.): p > 0.05, *0.05 > p > 0.01, and **p < 0.01. A/B and mA/B p=0.36, A/B and A-only p=0.003, A/B and mA/mB p=0.0005. (**C)** EMSAs comparing the effect of mutations in the A-box and B-box of ^His^tRNA gene upon TFIIIC binding. For each panel (mA/B, A/mB, A/B, and A-alone), a control without TFIIIC is performed (-), followed by increasing concentrations of TFIIIC as indicated (100, 250, 350 and 900 nM TFIIIC). The position for free DNA is indicated on the right as well as for the DNA/TFIIIC complex. (**D)** DNase I footprinting assay on the ^His^tRNA gene (120 bp, −44 to +76), labeled on the non-template strand with DS competitor. Lanes (left to right): M, marker; -, control (labeled DNA without DNase I); C, control (labeled non-template DNA with DNase I); labeled DNA with 4 or 8 pmol TFIIIC and DNase I; labeled DNA with TFIIIC (6 pmol) and TFIIIB (8 or 15 pmol); labeled DNA with TFIIIB alone and DNase I; M, marker. Second panel: same assay on ^His^tRNA gene with mutated A-box (mA/B). Yellow: A-box; Light Yellow: pseudo A-box; Light Blue: B-box; Brown: TATA-like site.

TFIIIC-bound lifetimes of 1.2 +/- 0.3 seconds (61.5 +/- 13.4 %) and 49 +/- 14 seconds (38.5 +/- 13.4 %) (Supplementary Figure 14A). Interestingly, live cell imaging of the yeast Pol III transcription machinery revealed similar residence times for TFIIIC τA (τA^Tfc4/τ131^ 33.4 s) and τB (τB^Tfc3/τ138^ 16.6 s) consistent with a model where τA stays bound longer compared to τB (33.4 s versus 16.6 s) because τA subsequently recruits TFIIIB (75).

In order to test the contribution of both A-box and B-box for binding TFIIIC, we repeated our smFRET experiments using three different mutant DNA constructs, which were informed from previous studies (11,76) or the τA-DNA cryo-EM structure obtained in this study: mA/B (mutations introduced only in the A-box), A-only (containing only the A-box) and mA/mB (mutations introduced only in the A- and B-box) (Figure 6A-B and Table S2 for sequence details). Simultaneously mutating both A-box and B-box or only containing the A-box, TFIIIC binding was almost completely abolished (Figure 6A-B, Supplementary Figure 14 C,D). This is in agreement with previous work (11,76) showing that the B-box is essential for TFIIIC binding. Unexpectedly, mutating four conserved nucleotides of the A-box, does not show a significant difference in TFIIIC binding dynamics (Figure 6A-B and Supplementary Figure 14B). Fitting the bound-lifetimes to a double-exponential function, we obtain TFIIIC-bound lifetimes of 0.8 +/- 0.1 seconds (80 +/- 10 %) and 54 +/- 31 seconds (20 +/- 10 %) (Supplementary Figure 14A). We verified these findings using electromobility shift assays where mutating the A-box also did not change TFIIIC-DNA binding significantly (Figure 6C). Overall, these results are consistent with a model in which the B-box is mainly responsible for TFIIIC engagement with its target site and the conserved A-box has functions other than contributing to the initial recruitment and retention of TFIIIC to DNA.

Having observed that mutating the major A-box does not significantly affect the TFIIIC DNA-bound lifetime (smFRET; Figure 6B and Supplementary Figure 14A-B) nor binding affinity (electromobility shift assays; Figure 6C) and our cryo-EM structures suggesting that τA mainly interacts with the DNA backbone rather than with base-specific interactions (Figure 4), we performed footprinting assays to investigate the contribution of the A-box in positioning the τA module (Figure 6D). In our footprinting experiments, we see protection of both B-box and A-box in the wildtype (WT) TFIIIC construct (A/B). In contrast, in the A-box mutant, the A-box footprint is almost lost. In order to investigate whether mutating the A-box has a downstream effect, we repeated our footprinting experiments in presence of TFIIIB (Figure 6D). We find that in A/B, A-box and B-box are still protected, but we observe a hypersensitivity to DNase I between the A- and B-box and between the A-box and the TATA-like element in the presence of TFIIIB. This suggests that TFIIIB induces bending of both DNA regions. In the mA/B mutant, although protection of the B-box is still visible, we observe no protection of the A-box and the TATA-like region, but also no TFIIIB-induced bending of the DNA, upon TFIIIC and TFIIIB addition. Taken together, our findings suggest that the A-box is less important for the initial binding of TFIIIC to DNA but that the A-box is important for correctly positioning τA which is critically required for the subsequent binding of TFIIIB.

## Discussion

The cryo-EM structure of the yeast TFIIIC complex bound to a ^His^tRNA gene offers new insights into the structural mechanism underlying TFIIIC mediated type II promoter recognition. Our study reveals the detailed interactions between the TFIIIC subcomplexes τA and τB and their conserved DNA target sites named A- and B-box. The large differences in binding affinities of τA and τB subcomplexes to the A- and B-box, respectively, is reflected in the number of protein-DNA interactions. Subcomplex τB interacts through numerous DNA backbone and DNA base-specific contacts with its high-affinity B-box target site (but also with DNA bases upstream the B-box) thereby almost completely enveloping the DNA. In contrast, τA shows only a few DNA interactions primarily with the DNA backbone and barely touches its low affinity A-box target site. We hypothesize that the A-box recognition is likely to occur through DNA shape readout.

Our cryo-EM data demonstrate the structural flexibility of the TFIIIC-DNA complex, and in particular the dynamic interaction of τA with its DNA-binding site compared to τB (Figure 7). Because of this flexibility, we could not obtain a complete 3D reconstruction of the entire TFIIIC complex, but only obtained independent reconstructions of the τA-DNA and τB-DNA complexes where the τA-DNA complex reconstruction showed lower resolution and greater variability in 2D classifications. Instead, cryo-EM particle pair distance mapping allowed us to determine the distance and relative orientation of both subcomplexes with a predominant inter-subcomplex distance of 168 Å, suggesting a flexible linkage between τA and τB that is essential for TFIIIC’s function in promoter recognition. The flexible and dynamic nature of τA is likely crucial for its role in locating the A-box on tRNA genes and recruiting other transcription factors such as TFIIIB.

**Figure 7.**
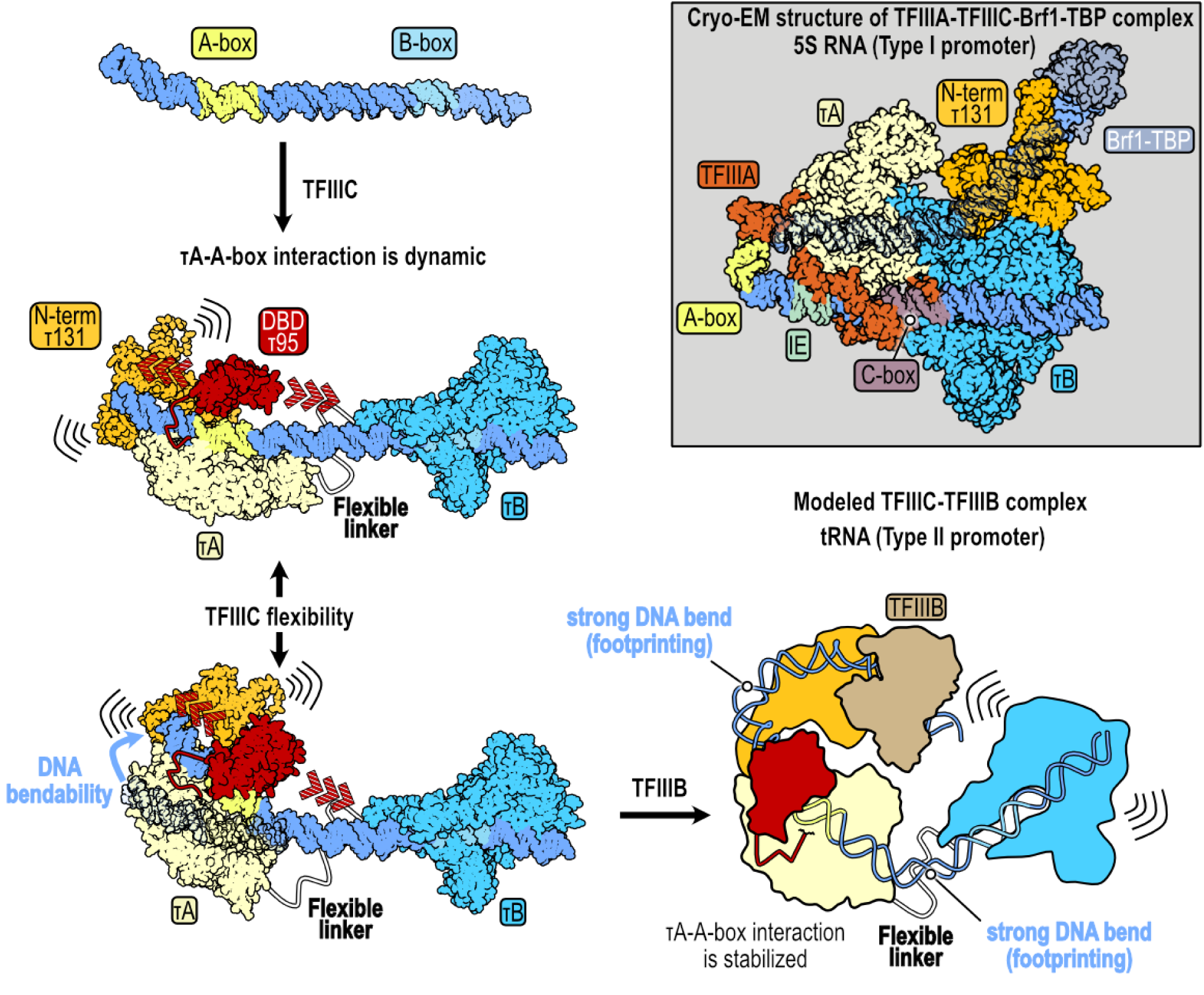
Model of TFIIIC-TFIIIB assembly on type II promoter. The ^His^tRNA, containing A- and B-box elements spaced by 32 nucleotides, binds to TFIIIC. TFIIIC stably interacts with the B-box via the τB subcomplex, while the τA subcomplex binds dynamically to the upstream region through DNA-backbone contacts. Additionally, the putative DNA-binding domain (DBD) of τ95 and the N-terminal TPR domain repeats of τ131 exhibit high flexibility. The flexible linker connecting τA and τB allows TFIIIC to adapt to the highly flexible DNA, which is prone to bending between these subcomplexes. Upon recruitment of TFIIIB, mediated by the flexible N-terminal TPR domain repeats of τ131 to the TATA-like region, the newly formed TFIIIC-TFIIIB complex undergoes a conformational change. This change induces bending of the DNA between the TATA-like region and the A-box, as well as between the A-box and the B-box, resembling the TFIIIA-TFIIIC-Brf1-TBP complex observed in the type I promoter.

Comparative structural analysis between yeast and human TFIIIC revealed both conserved and distinct features. Despite only 18% sequence identity between yeast and human τ138, the spatial configuration of the WH domains around the B-box is strikingly similar, emphasizing the evolutionary conservation of these critical DNA-binding regions when bound to the conserved B-box DNA. In contrast, we could not observe a DNA-bound τA complex in human TFIIIC, although cryo-EM particle pair distance mapping confirmed that also human τA and τB are stably, but flexibly, linked subcomplexes (32).

Comparison with yeast TFIIIA-TFIIIC and TFIIIA-TFIIIC-Brf1-TBP bound to type I promoter DNA (33) provides additional insights. The presence of TFIIIA dramatically changes the interaction of τA and τB with their target sites. In the TFIIIA-TFIIIC-DNA complex, TFIIIA binds prior to TFIIIC to the IC element/C-box of the type I promoter thereby considerably changing the way TFIIIC interacts with the DNA (Supplementary Figure S9). In the next step, the addition of TBP-Brf1 leads to a dramatic bending of type I promoter DNA that now undergoes a 180° turn together with the rearrangement and change of conformation of the N-terminal TPR domain of τ131 that also becomes more ordered (Figure 7). In the absence of a corresponding structure of a TFIIIC-Brf-TBP complex bound to a type II promoter, we can only speculate whether this complex bound to type II promoter undergoes similar dramatic conformational changes and adopts a compact conformation like the TFIIIA-TFIIIC-Brf1-TBP-DNA complex. However, two observations suggest that this could be the case. First, footprinting results clearly indicate strong increased DNA backbone accessibility to DNase I upon the addition of TFIIIB (Figure 6D) that were also observed by others (77,78) indicating DNA bendability between the A- and B-box and between the A-box and the TATA-like element. Second, a direct interaction between TBP and τB subunit τ60 (30, 34) suggests that TFIIIB and τB are in close proximity as part of a compact complex. Single-molecule experiments and EMSA experiments suggest that τA binding to the A-box play only a minor role in TFIIIC residence times to type II promoter DNA compared to τB and the B-box. Instead, the recognition of the A-box by the τA subcomplex and the deformability of DNA upstream and downstream of the A-box play critical roles in the recruitment of TFIIIB and Pol III and assigns τA and the A-box important roles in Pol III pre-initiation complex assembly.

## Data Availability

Cryo-EM maps of the yeast TFIIIC subcomplexes τA and τB bound to DNA have been deposited to the Electron Microscopy Data Base (EMDB) under accession codes EMD-51231 (τA-DNA monomer) and EMD-51228 (τB-DNA monomer). Atomic coordinates of the two subcomplexes bound to DNA have been deposited to the Protein Data Bank under accession codes: PDB ID 9GCK (τA-DNA monomer) and PDB ID 9GC3 (τB-DNA monomer).

## Supplementary Data

Supplementary Data are available at NAR Online.

## Acknowledgements

We acknowledge support by K. Lapouge (EMBL Protein Expression and Purification Core Facility), M. Rettel (EMBL Proteomics Core Facility), and Joseph Barto (EMBL Cryo-EM Service Platform). We also thank H. Fung and J. Weidenhausen for support and discussion. W.S.-D. acknowledges support from the EMBL International PhD program. W.S.-D., A.C., F.B., L.H., T.H., O.D., S.E., and C.W.M. acknowledge support by EMBL.

