## Supplementary Data for "Structural and kinetic insights into tRNA promoter engagement by yeast general transcription factor TFIIIC"

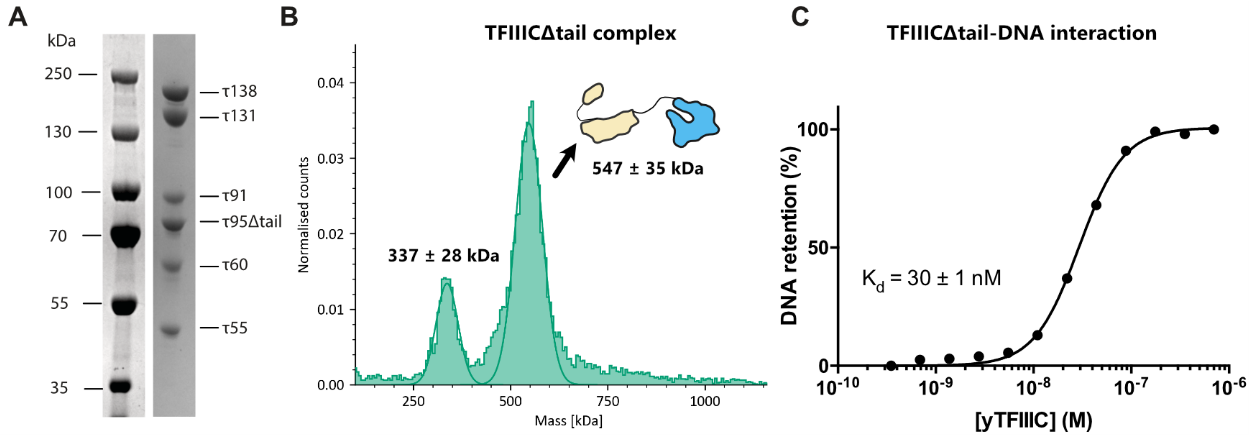


**Supplementary Figure S1. Analysis of the structural integrity and DNA interaction of the TFIIICΔtail complex**. **(A)** SDS-PAGE illustrating the homogeneity of the sample. The six subunits that form the TFIIICΔtail complex are labeled on the right. **(B)** Mass photometry experiment reveals an intact TFIIICΔtail complex. The graph shows a primary peak representing the full TFIIICΔtail complex, with an expected molecular mass of 547 ± 35 kDa, and a smaller peak at 337 ± 28 kDa, which likely corresponds to a disassembled subcomplex. **(C)** Filter binding assay. TFIIICΔtail was titrated against the radiolabeled 85 bp yeast $^{His}$tRNA gene. The binding data were fitted to a Hill equation with a fixed Hill coefficient of 1. The dissociation constant ($K_d$) was estimated to be 30 nM, with a standard error of 1 nM, N = 1.

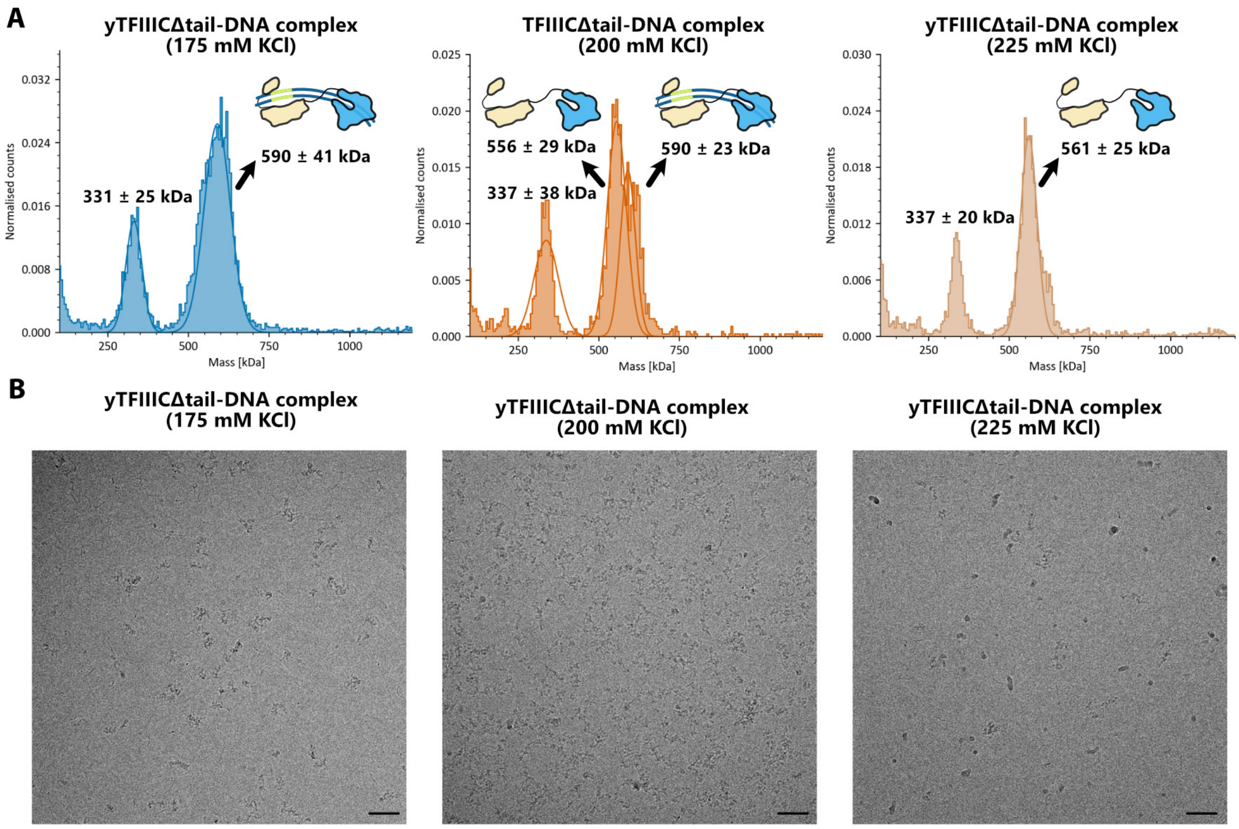


**Supplementary Figure S2. Mass photometry and cryo-EM assessment of yTFIIICΔtail-DNA complex stability under various ionic conditions. (A)** Mass photometry analysis illustrates the stability of the TFIIICΔtail-DNA complex under diverse salt concentrations. Each plot is labeled with the salt concentration examined. **(B)** Cryo-electron microscopy micrographs demonstrate the complex under the same ionic conditions as those assessed by mass photometry, featuring a scale bar corresponding to 500 Å. Images were captured at a magnification of 92,000x and a pixel size of 1.566 Å.

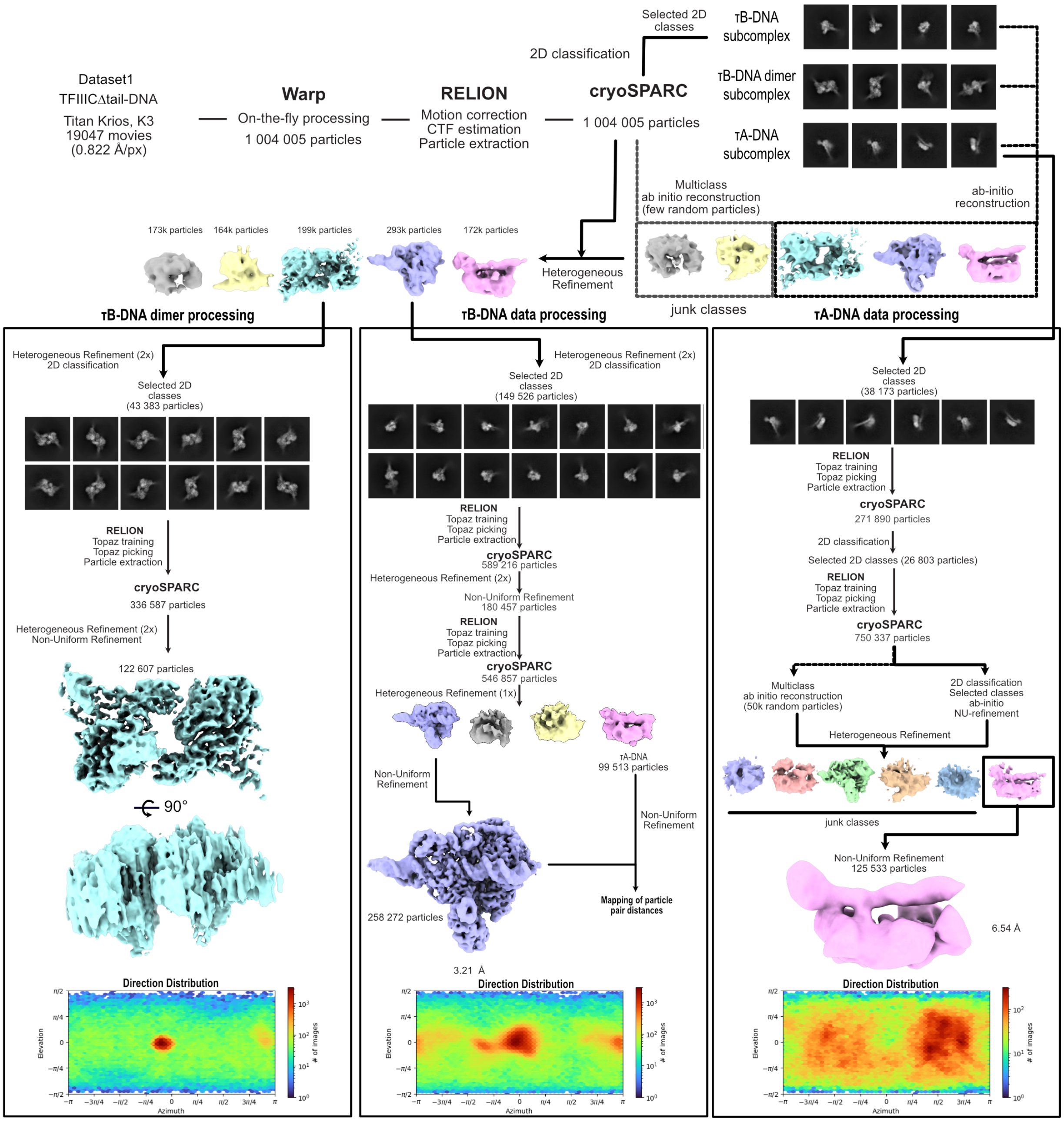


**Supplementary Figure S3. Workflow for the first cryo-EM dataset of the TFIIICΔtail-DNA complex.** This dataset was initially classified into three categories: τB-DNA dimer (left), τB monomer (middle), and τA-DNA (right) subcomplexes. Each classification underwent particle training and picking using TOPAZ, followed by several rounds of heterogeneous refinement. The τB-DNA dimer class, utilizing particles from the TOPAZ picking step, after two rounds of heterogeneous refinement, yielded a map containing 122,607 particles with preferred orientation. The τB-DNA monomer, also after two TOPAZ rounds of training/picking, resulted in a map of 258,272 particles with a resolution of 3.21 Å. Similarly, the τA-DNA subcomplex, following the same TOPAZ and refinement steps, produced a map from 125,533 particles at a resolution of 6.54 Å. During the τB-DNA monomer data processing, τA-DNA particles picked in the second TOPAZ round were used for mapping particle pair distances. Particle distribution for each class is displayed in the lower section of their respective data processing pipeline.

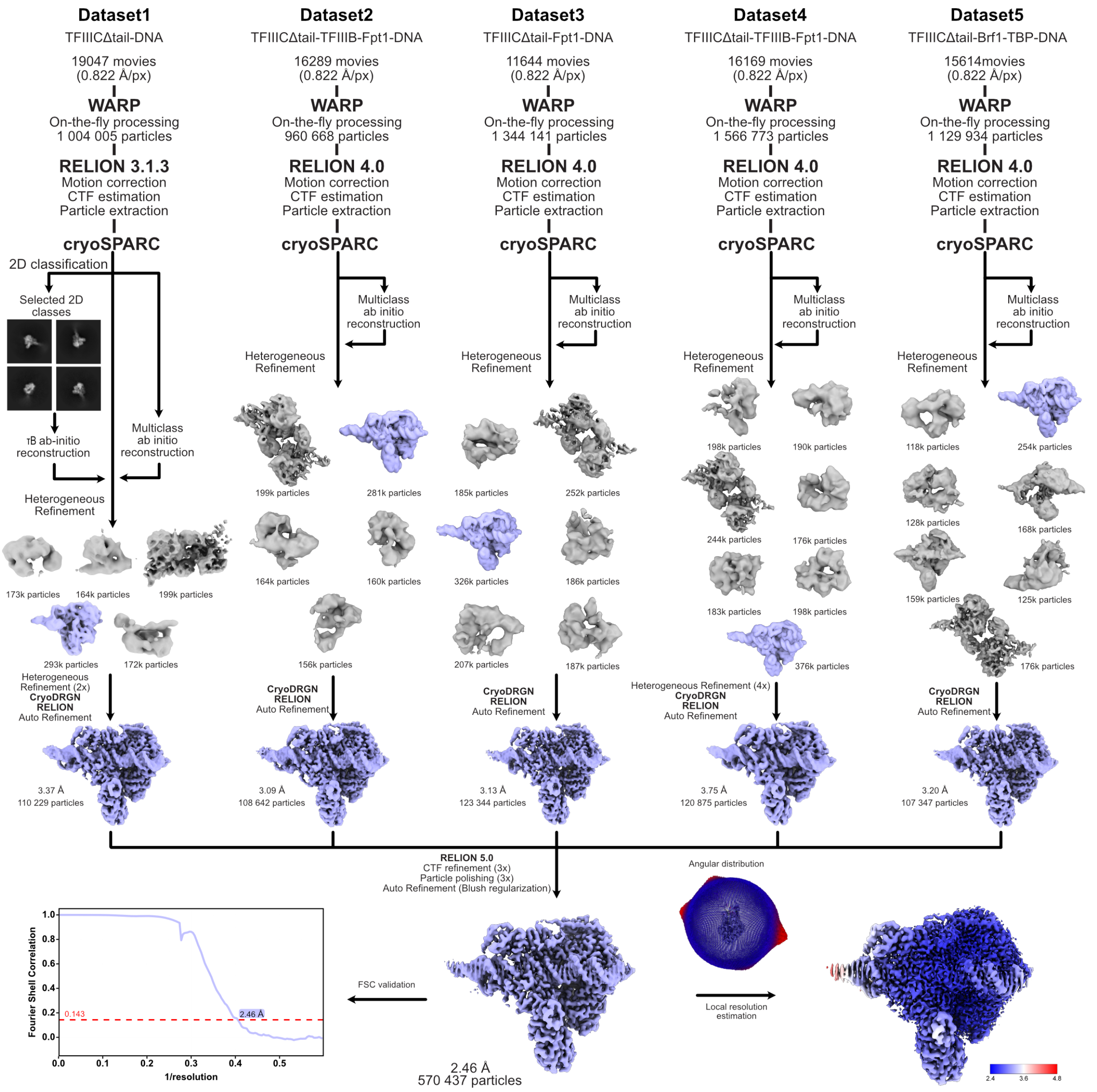


**Supplementary Figure S4. General workflow for processing all cryo-EM datasets focusing on the τB-DNA subcomplex.** A comprehensive processing workflow for the τB-DNA subcomplex across five cryo-EM datasets is shown. Initial on-the-fly data processing was performed in WARP, which included particle picking for each dataset. Following extraction in Relion, these particles were imported to cryoSPARC. An ab initio reconstruction step, targeting particles featuring the τB-DNA subcomplex, was carried out, followed by a multiclass *ab initio* reconstruction to generate decoy volumes to help sort out junk particles. Several rounds of heterogeneous refinement were conducted to obtain a homogeneous τB-DNA particle population. Before combining particles from all five datasets into Relion5.0, cryoDRGN was used to further address remaining heterogeneity. The merged particle set, 570,437 particles, yielded a map at 3 Å resolution. Further refinement through three rounds of CTF refinement and particle polishing improved the resolution to 2.46 Å. The FSC curve of this refined map is shown at the bottom left, and the angular distribution of the particles, alongside the local resolution estimation map, is displayed at the bottom right. Additionally, representative 2D class averages, 3D classifications from heterogeneous refinement, Auto refinement maps of τB-DNA, and junk volumes are also illustrated.

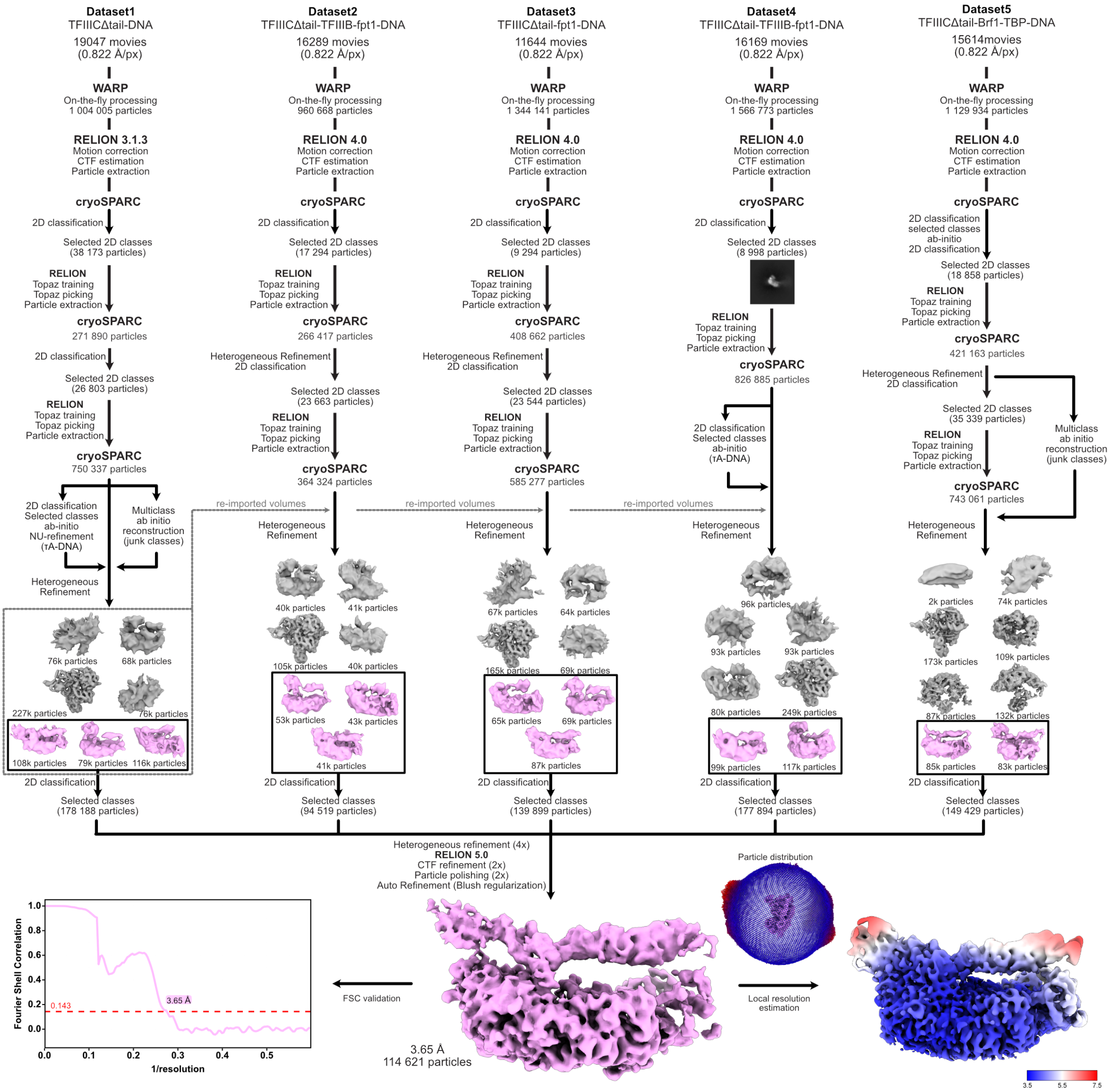


**Supplementary Figure S5: General workflow for processing all cryo-EM datasets focused on the τA-DNA subcomplex**. The initial steps included on-the-fly processing and particle picking in WARP, followed by particle extraction in Relion and further processing in cryoSPARC. To increase the initial particle number from WARP, two rounds of TOPAZ training and picking were conducted for each dataset, except for dataset 4, which underwent only one round. Initial ab initio reconstructions of the τA-DNA subcomplex and decoy volumes were generated during the processing of dataset 1. These volumes were then imported into datasets 2-4 for subsequent heterogeneous refinement. For dataset 5, new *ab initio* τA-DNA and decoy maps were reconstructed. After the final heterogeneous refinement of each dataset, particles were further cleaned using 2D classification, discarding only obvious junk classes before merging. The merged particles underwent four rounds of heterogeneous refinement, resulting in a map containing 114,321 particles. These were then re-imported into Relion5.0, where they reached a resolution of 3.65 Å after two rounds of CTF refinement and particle polishing. The figure also displays the FSC curve (cutoff = 0.143) of the final volume (left), the angular distribution of the particles, and the local resolution estimation map (both at bottom right). Additionally, representative 2D class averages, 3D classifications from heterogeneous refinement, and auto-refinement maps of τA-DNA, along with decoy volumes, are also included.

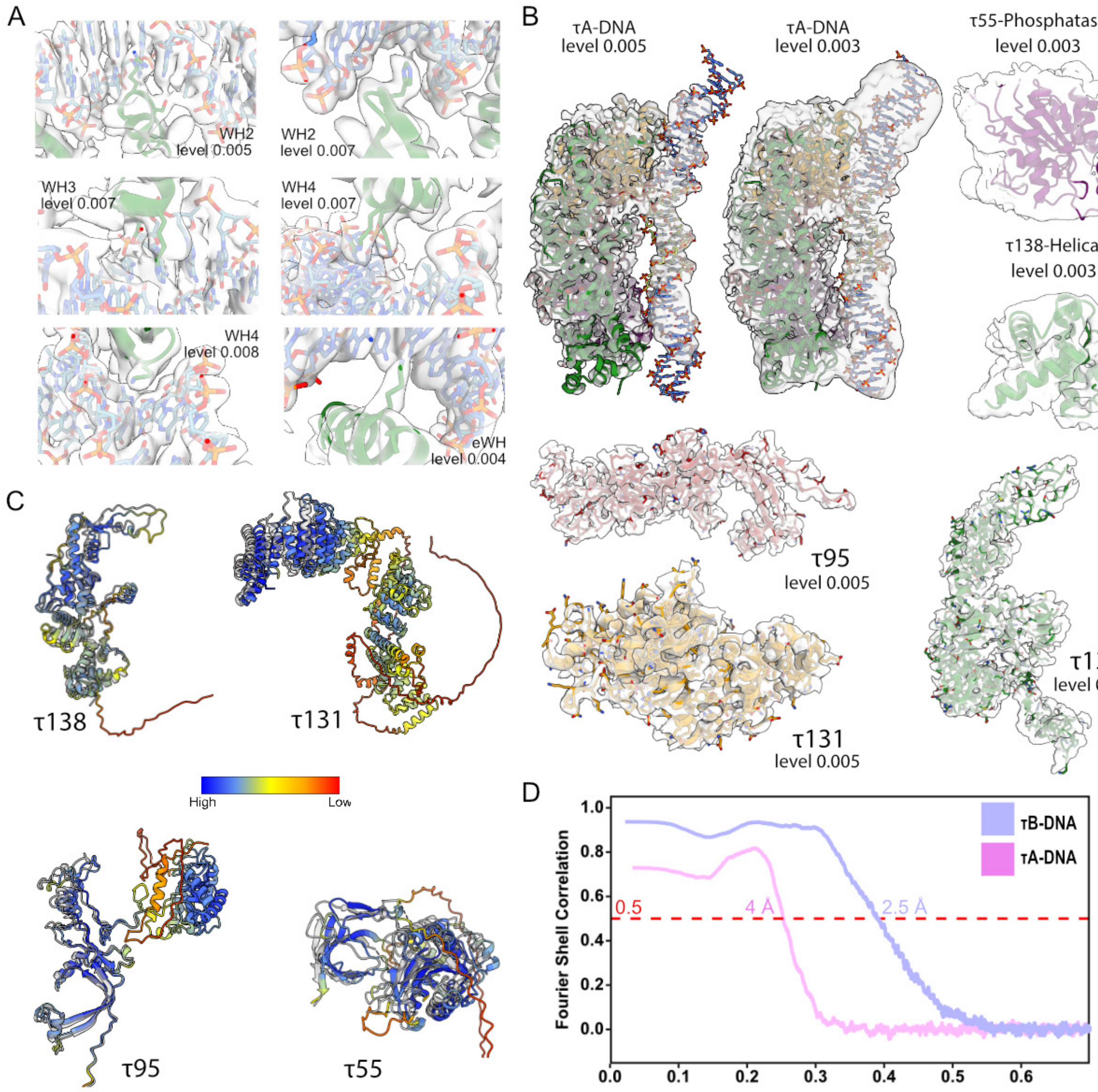


**Supplementary Figure S6: Alphafold predictions and quality of map densities. (A)** Cryo-EM densities with refined models showing side chains of amino acids from τ138 involved in base-specific contact with DNA and B-box. **(B)** Exemplary models of τA-DNA, 45bp $^{his}$tRNA DNA, τ55-Phosphatase, τ138-helical, τ138, τ131, and τ95, each distinctly colored and labeled, with threshold levels indicated. **(C)** Alphafold predictions for each τA subunit superimposed with built models in grey. The multimer prediction was separated into its individual components for clarity **(D)** FSC curves for τA-DNA and τB-DNA maps and models, with the FSC = 0.5 threshold marked by a dashed red line.

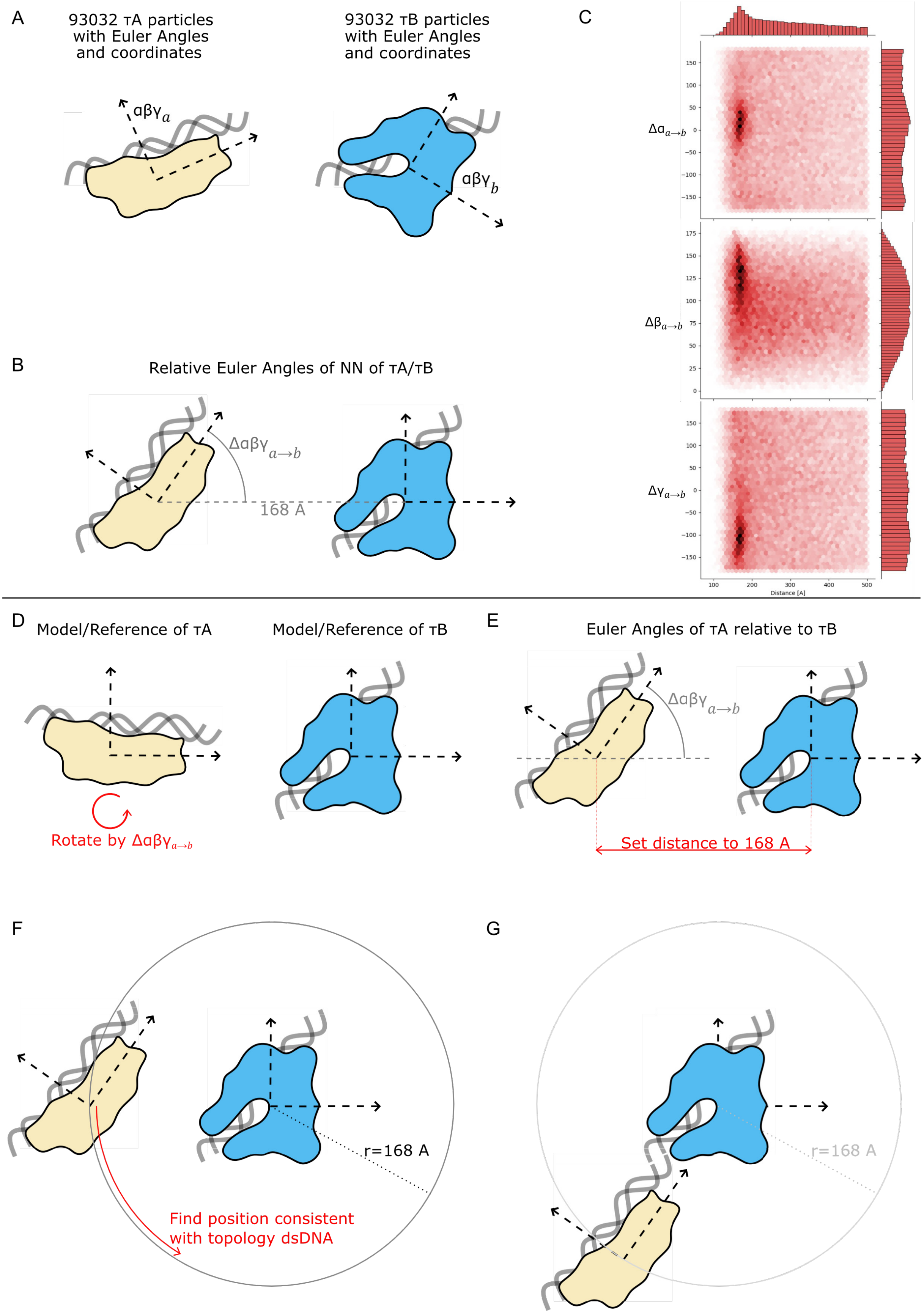


**Supplementary Figure S7. Relative orientations of τA-DNA and τB-DNA subcomplexes. (A**) The 2D coordinates and the Euler angles αβγ of the τA and τB particles were obtained during the individual picking and refinement steps. For clarity τA and τB particles and angles are illustrated in 2D. **(B)** Based on nearest neighbor proximity search, pairs of τA and τB particles were identified and assigned. The distance between the τA and τB particles in the XY-plane, as well as the orientation of τA relative to τB, were calculated from their respective positions and Euler angles. **(C)** The three relative Euler angles $\Delta\alpha\beta\gamma_{a\rightarrow e}$ of each τA/τB complex were plotted against their distance. The distance distribution exhibits a peak at 168 Å. The Euler angles show enrichment around Δα ≈ 15°, Δβ ≈ 128°, and Δγ ≈ -103°. This enriched population was observed for particle pairs separated by 150 Å to 190 Å. The structure of the τA-τB-DNA complex was modeled based on individually refined volume maps. τA was positioned and oriented relative to a τB. **(D)** Initially, τA was rotated by $\Delta\alpha\beta\gamma_{a\rightarrow b}$ to obtain the correct relative orientation. The chosen Euler angles for $\Delta\alpha\beta\gamma_{a\rightarrow e}$ correspond to the values Δα=15°, Δβ=128°, and Δγ=-103°. **(E)** Subsequently, both models were placed at the most abundant distance value of 168 Å. **(F)** This configuration constrained the degree of freedom of τA to translational movement on the surface of a sphere, with τB at the center and a radius of 168 Å. **(G)** The remaining degree freedom could be resolved by the topology of the DNA.

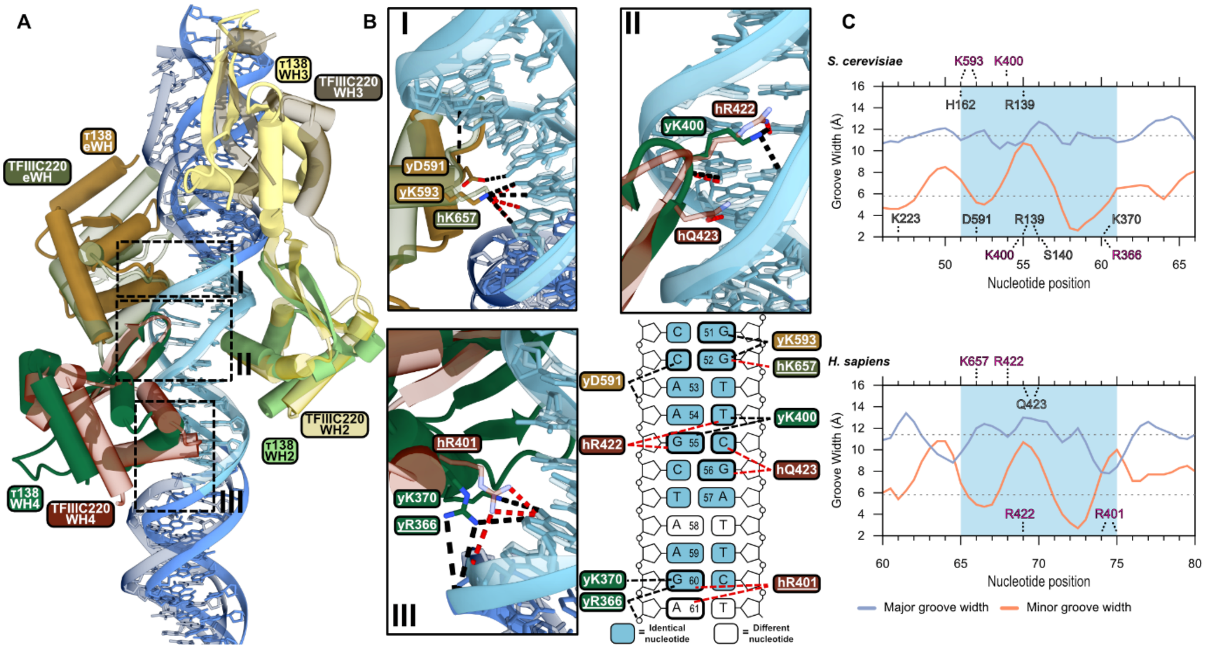


**Supplementary Figure S8. Comparative analysis of winged-helix (WH) domains binding to tRNA genes in yeast and human τB subcomplexes. (A)** WH domains from both yeast and human complexes have been superimposed using the B-box promoter region DNA sequence in ChimeraX to illustrate structural similarities and differences. **(B)** Detailed views of each comparison are provided to highlight the amino acids that form hydrogen bonds in a base-specific manner with the DNA. In these visualizations, human WH domains and its tRNA associated gene are depicted as transparent models to differentiate them from their yeast counterparts. The schematic illustrates the base-specific contacts of the highlighted amino acids within the B-box of the yeast $^{His}$tRNA gene. Red dotted lines denote interactions specific to the human sequence, while black dotted lines represent interactions in the yeast sequence. The sequence displayed belongs to the B-box region of the yeast $^{His}$tRNA gene. Nucleotides highlighted in light blue indicate positions that are conserved between yeast and human B-box sequences, whereas nucleotides shown in white represent divergent positions between the two species. **(C)** DNA groove width analysis of the *S. cerevisiae* and *H. sapiens* τB-DNA complexes. Both analyses were computed with the Curves+ software (1). Residues that form base-specific contacts with nucleotides of the non-template and template DNA strand are shown at the top and bottom of the plots, respectively. Those residues that are identical or highly conserved amongst a broad range of eukaryotes (see Seifert et al. (2)) are colored in pink. The light-blue shading marks the location of the B-box.

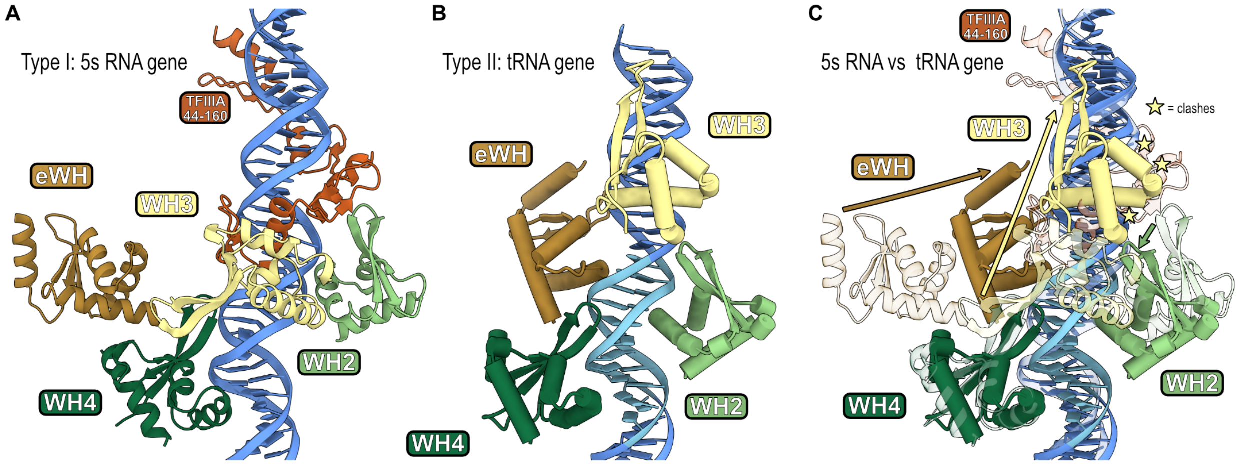


**Supplementary Figure S9. Comparative analysis of yeast τB subcomplexes bound to type I and type II Pol III promoters. (A)** The τB subcomplex bound to a 5S RNA gene is presented as a ribbon model where only the WH domains are visible. The TFIIIA region from amino acids 44 to 160 is shown. The rest of the TFIIIA-TFIIIC-TFIIIB complex was omitted for clarity (PDB ID: 8FFZ). **(B)** Cryo-EM model from this study of the yeast τB subcomplex bound to a tRNA gene, depicted as a cylinder model with a focus on the WH domains. Other domains and loops are hidden to enhance visual clarity; here, the B-Box promoter is highlighted in light blue. **(C)** Superimposition of both τB subcomplexes from panels A and B, using the τ91 and τ60 subunits as reference points. In this comparison, the cryo-EM model of τB bound to the type I promoter is rendered transparent to delineate differences in the positioning of the WH domains, which are indicated by arrows. These arrows are color-coded to correspond with each WH domain, illustrating the spatial variances between the two gene types. The yellow stars indicate clashes between WH3 and TFIIIA.

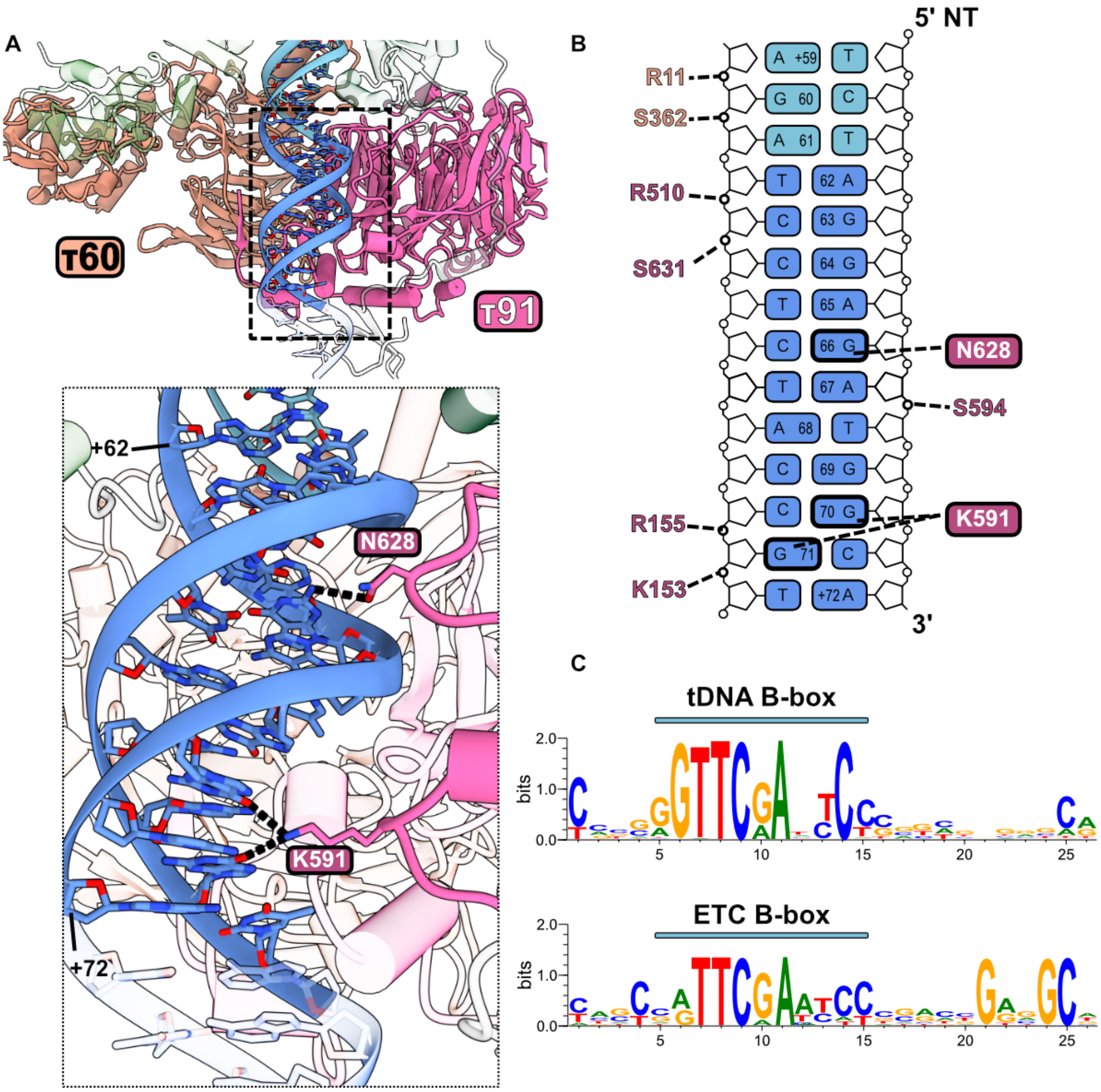


**Supplementary Figure S10. Base-specific interactions of τ91 with the region downstream of the B-box. (A)** Interactions of subunits τ91 and τ60 with DNA. The inset provides a close-up view of τ91, highlighting the amino acids involved in base-specific interactions, each marked within colored squares. **(B)** Schematic representation of all polar interactions formed between subunits τ91 and τ60 with DNA. **(C)** Sequence logos of the B-box and nearby regions from all yeast tRNA genes (top) and from all ETC sequences (bottom).

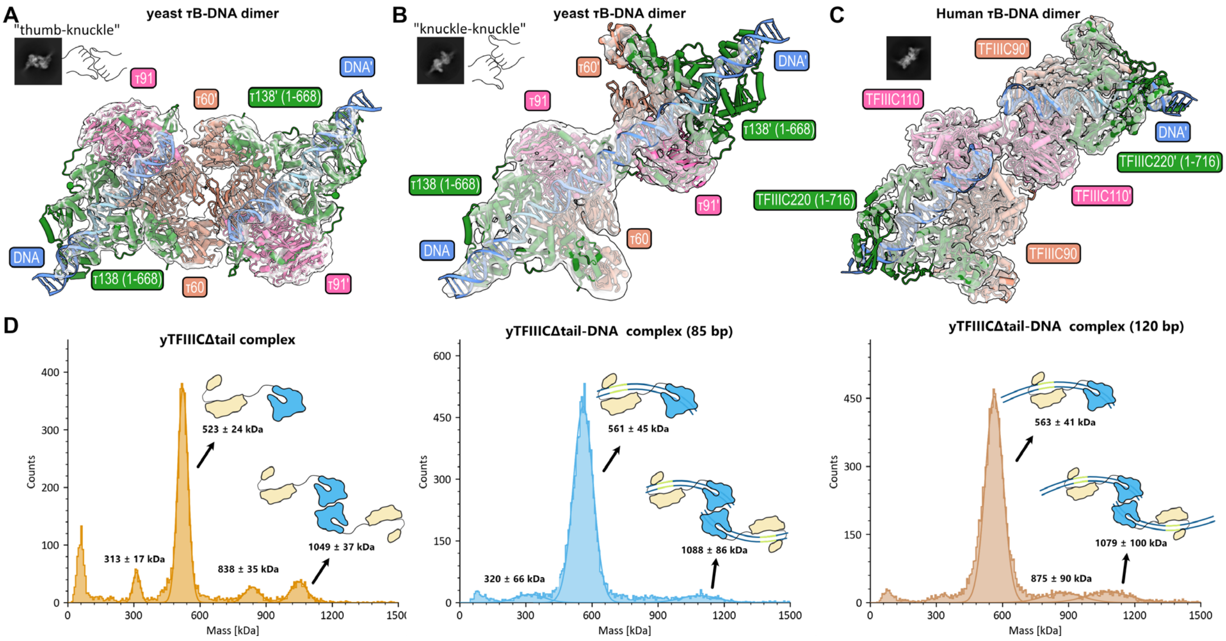


**Supplementary Figure S11. Differences in dimerization between yeast and human τB-DNA subcomplexes. (A)** In yeast, two distinct forms of τB-DNA complex dimerization are observed. The first form, referred to as "thumb-knuckle," involves interactions between the τ60 subunits (equivalent to TFIIIC90 in humans) of each monomer. **(B)** The second form, termed "knuckle-knuckle," is mediated through the τ91 subunits (equivalent to TFIIIC110 in humans) and the "latch" domain within τ138. **(C)** In contrast, the dimerization of the human τB-DNA complex predominantly involves the TFIIIC110 subunits without engagement from other subunits. Representative 2D classes for each dimer type are displayed. **(D)** Mass photometry analysis of yTFIIICΔtail and yTFIIICΔtail-DNA complex dimerization at a concentration of 100 nM: (Left) Mass distribution histograms for the yTFIIICΔtail complex without DNA show a main peak at 523 ± 24 kDa (monomer), with higher molecular weight peaks at 838 ± 35 kDa and 1049 ± 37 kDa, indicating dimer formation. (Middle) Mass distribution histograms for the yTFIIICΔtail complex with 85 bp DNA show a main peak at 561 ± 45 kDa (monomer-DNA complex), with a significant higher molecular weight peak at 1088 ± 86 kDa (dimer-DNA complex). (Right) Mass distribution histograms for the yTFIIICΔ593 complex with 120 bp DNA exhibit a main peak at 563 ± 41 kDa (monomer-DNA complex), with higher peaks at 875 ± 90 kDa and 1079 ± 100 kDa (dimer-DNA complex). Accompanying schematic diagrams next to each analyzed peak illustrate the TFIIICΔtail and TFIIICΔ593-tail complexes in monomer and dimer forms, with molecular weights specified below each diagram. The consistent first peak across all conditions, corresponding to a protein with a molecular weight of 313-320 kDa, suggests a potential disassembly of the full TFIIIC complex into one of its subcomplexes.

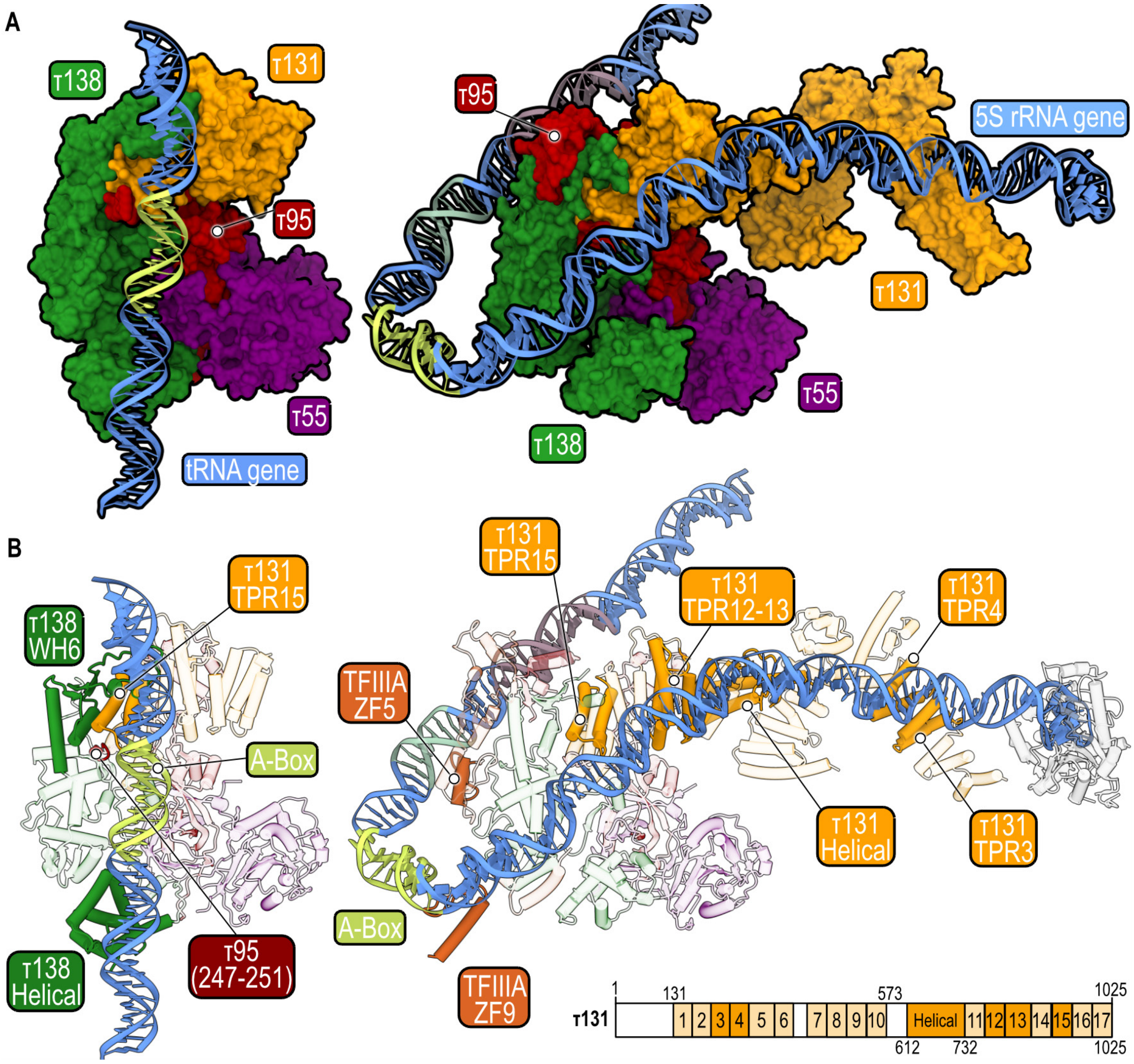


**Supplementary Figure S12: Comparative analysis of yeast τA subcomplex bound to type I and type II Pol III genes**. **(A)** Cryo-EM models of the τA subcomplex bound to type I and type II promoters. Left: τA subcomplex bound to a tRNA gene. Right: τA subcomplex bound to the 5S rRNA gene. Note that TFIIIA and Brf1-TBP are not depicted. **(B)** Domains and regions of different subunits involved in DNA binding are shown in both τA-DNA complexes. Left: DNA-binding domains for the tRNA gene. Right: DNA-binding domains for the 5S rRNA gene. The domain architecture of the τ131 subunit is depicted, illustrating all the TPR domain repeats and the helical domain along the subunit. The domains that interact with DNA are shown in full color, while those that do not interact are shown in transparent. Key DNA regions are highlighted: A-box in yellow, internal element (IE) in light green, and C-box in grey.

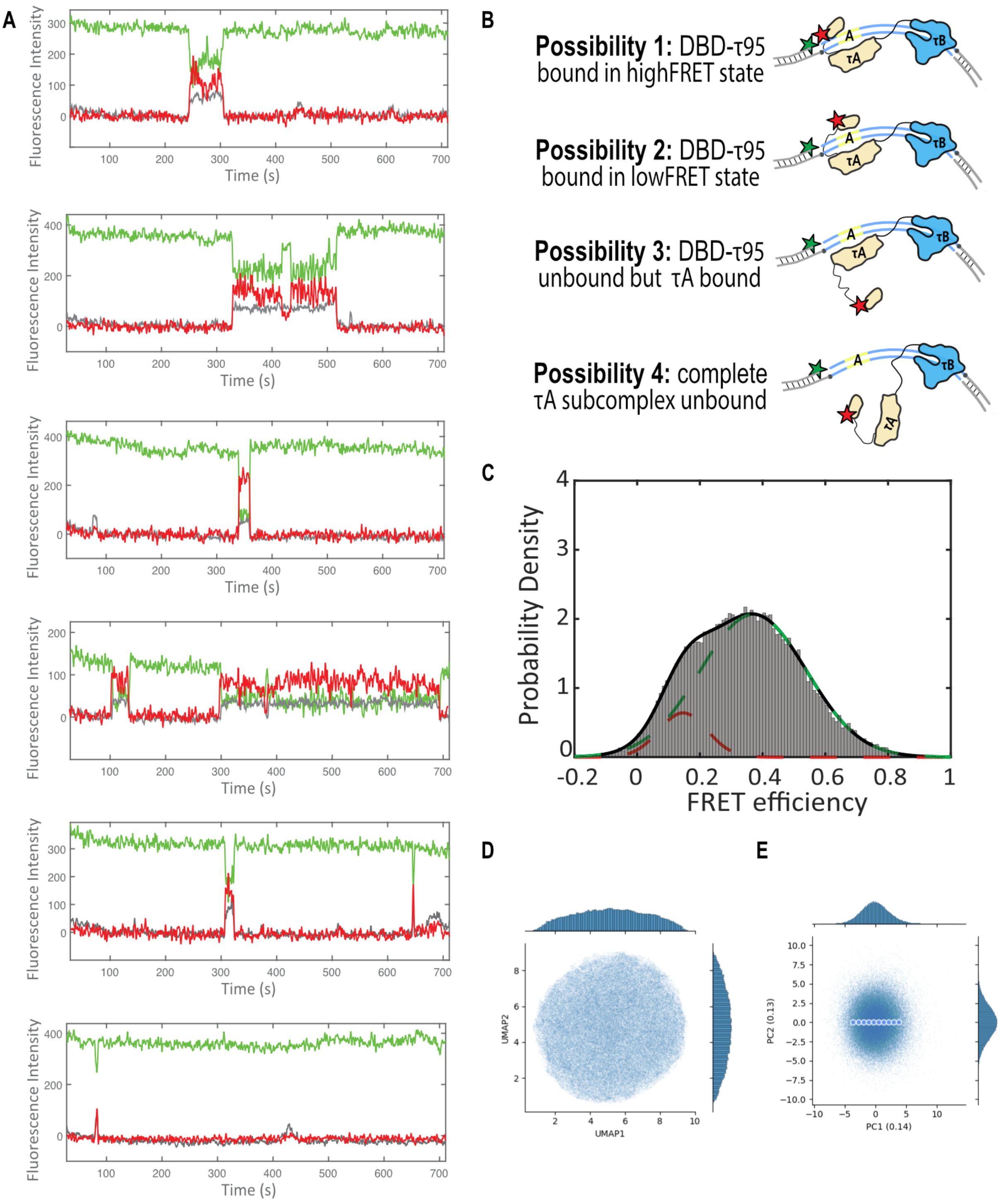


**Supplementary Figure S13: Analysis of TFIIIC binding and τA-DNA conformational dynamics. (A)** Representative single-molecule traces of TFIIIC-Cy5 binding to A/B DNA**. (B)** Schematic representation of possible TFIIIC bound states. **(C)** FRET efficiency distribution of TFIIIC-Cy5 bound to A/B DNA-Cy3 (same data as in Figure 5D) fitted to a two Gaussian function. $\mu 1 = 0.15 +/- 0.07$, $\mu 2 = 0.37 +/- 0.17$. **(D)** UMAP visualization of the 8-dimensional latent space representation learned by cryoDRGN for τA-DNA particle images. **(E)** Principal Component Analysis (PCA) projection of the 8-dimensional latent encodings from cryoDRGN, with blue points corresponding to the trajectory across the PC1. Four maps from PC1 were used in Figure 5C.

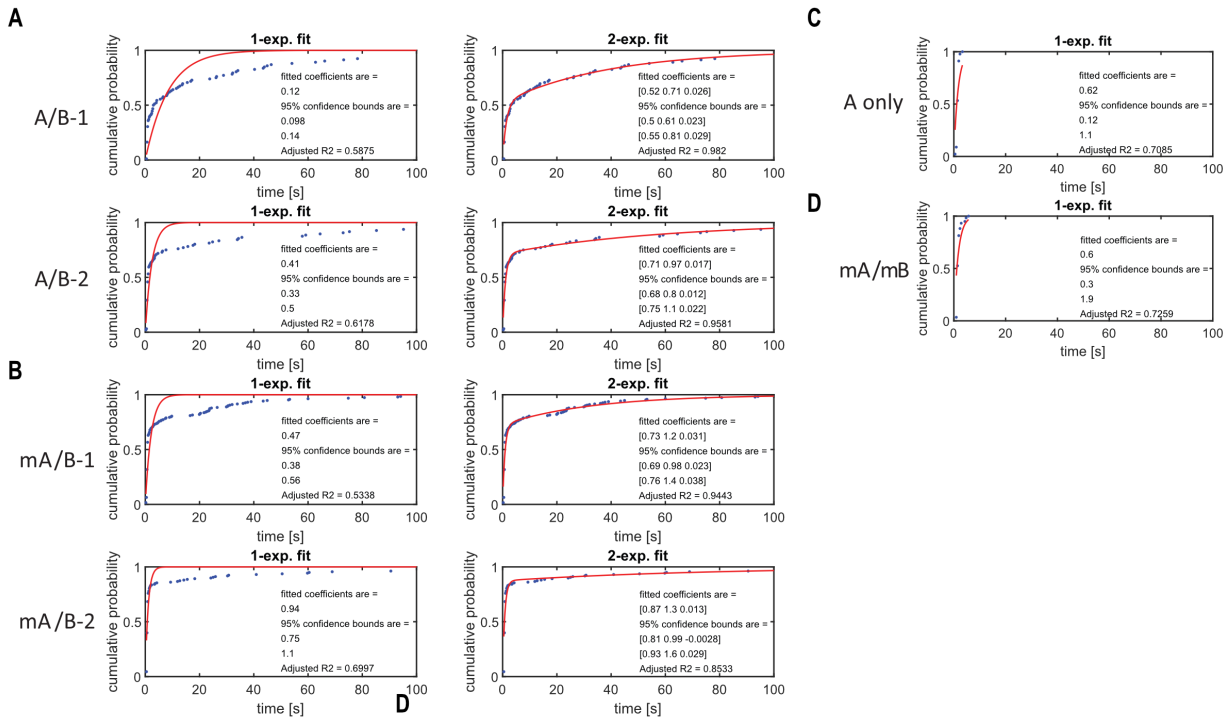


**Supplementary Figure S14**. Single and double exponential fittings of TFIIIC-bound lifetimes. For single exponential fit, the coefficient corresponds to the off-rate (sec-1). For the double exponential fit, the three coefficients correspond to population of first state, first and second off-rates (both in sec-1). **(A)** TFIIIC-bound lifetimes to A/B DNA, number of dwells analysed (n) = 92, 96. **(B)** mA/B DNA, (n) = 189, 158. **(C)** A only, n = 45. **(D)** mA/mB DNA, n = 59.

### Table S1a. Cryo-EM data collection

| | Dataset 1<br>TFIIICΔtail-DNA | Dataset 2<br>TFIIICΔtail-TFIIIB-Fpt1-DNA | Dataset 3<br>TFIIICΔtail-Fpt1-DNA | Dataset 4<br>TFIIICΔtail-TFIIIB-Fpt1-DNA | Dataset 5<br>TFIIICΔtail-Brf1-TBP-DNA |
|---|---|---|---|---|---|
| **Data collection** | | | | | |
| Magnification | 105,000 | 105,000 | 105,000 | 105,00 | 105,00 |
| Voltage (kV) | 300 | 300 | 300 | 300 | 300 |
| Electron exposure (e–/Å$^2$) | 39.6 | 43.6 | 43.2 | 43.2 | 44.4 |
| Defocus range (μm) | 0.7-1.7 | 0.7-1.7 | 0.7-1.7 | 0.7-1.7 | 0.7-1.7 |
| Pixel size (Å) | 0.822 | 0.822 | 0.822 | 0.822 | 0.822 |
| Symmetry imposed | C1 | C1 | C1 | C1 | C1 |
| Micrographs (no.) | 19,047 | 16,289 | 11,644 | 16,169 | 15,614 |
| Initial particle images (no.) | 1,004,005 | 960,668 | 1,344,141 | 1,566,773 | 1,129,934 |
| Final τB-DNA particle images before data merging (no.) | 110,229 | 108,642 | 123,344 | 120,875 | 107,347 |
| Final τA-DNA particle images before data merging (no.) | 178,188 | 94,519 | 139,899 | 177,894 | 149,429 |

### Table S1b. Cryo-EM refinement and validation statistics

| | τB-DNA | τA-DNA |
|---|---|---|
| Initial particle images after data merging (no.) | 570,437 | 739,929 |
| Final particle images after merging (no.) | 570,437 | 114,621 |
| Map resolution (Å) | 2.46 | 3.65 |
| FSC threshold | 0.143 | 0.143 |
| Map resolution range (Å) | 2.39-5.56 | 3.51-7.28 |
| **Refinement** | | |
| Initial model used | Alphafold multimer prediction (v 2.3.0) | Alphafold multimer prediction (v 2.3.0) |
| Model resolution (Å) | 2.5 | 4 |
| FSC threshold | 0.5 | 0.5 |
| Map sharpening B factor (Å2) | -66.84 | -105 |
| **Model composition** | | |
| Non-hydrogen atoms | 15049 | 12301 |
| Protein/Nucleotide residues | 1673 / 80 | 1287 / 90 |
| Ligands | - | - |
| **B factors** (Å2) | | |
| Protein (min/max/mean) | 23.05/613.48/98.76 | 19.66/440.00/245.56 |
| Nucleotide (min/max/mean) | 24.79/617.20/144.81 | 142.73/440.00/404.64 |
| **R.m.s. deviations** | | |
| Bond lengths (Å) | 0.013 | 0.015 |
| Bond angles (°) | 1.641 | 1.942 |
| **Validation** | | |
| MolProbity score | 0.89 | 1.91 |
| Clashscore | 1.26 | 10.65 |
| Poor rotamers (%) | 0.00 | 0.77 |
| **Ramachandran plot** | | |
| Favored (%) | 97.84 | 94.74 |
| Allowed (%) | 2.16 | 5.03 |
| Disallowed (%) | 0.0 | 0.24 |
| **Rama-Z score** | | |
| Whole | -0.68(0.19) | -2.69(0.21) |
| Helix | -0.97(0.23) | -2.43(0.16) |
| Sheet | 0.12(0.23) | -0.46(0.56) |
| Loop | -0.36(0.21) | -1.23(0.23) |

**Table S2. Adapter and tRNA gene oligonucleotides used for single molecule fluorescence microscopy experiments and EMSA**

| Oligonucleotide | DNA sequence |
|---|---|
| p0074-Cy3 | 5'- /5Cy3/GG TGT ATG TAA TTG GAG TGG TT -3' |
| p0141 | 5'-GGT GTG TTG GTT GGG TGG TGG TAG T TGTGG AATTG TGAGC GGATA A -3’ |
| p0109-biotin | 5'-5BiotinTEG/ TTA TCC GCT CAC AAT TCC ACA -3’ |
| A/B | 5'-TCCATTGAAAAGTCGCCATCTTAGTATAGTGGTTAGTACACATCGTTGTGGCCGATGAAACCCTGGTTCGATTCTAGGAGATGGCATTTT -3’ |
| mA/B * | 5’-TCCATTGAAAAGTCGCCATCTAACTATTGTCGTTAGTACACATCGTTGTGGCCGATGAAACCCTGGTTCGATTCTAGGAGATGGCATTTT -3’ |
| mA/mB * | 5’-TCCATTGAAAAGTCGCCATCTAACTATTGTCGTTAGTACACATCGTTGTGGCCGATGAAACCCTGGACGCATTCTAGGAGATGGCATTTT -3’ |
| A only | 5’-TCCATTGAAAAGTCGCCATCTTAGTATAGTGGTTAGTACACATCGTTGTG-3’ |

* The mutated nucleotides are underlined.
